# nELISA: A high-throughput, high-plex platform enables quantitative profiling of the inflammatory secretome

**DOI:** 10.1101/2023.04.17.535914

**Authors:** Milad Dagher, Grant Ongo, Nathaniel Robichaud, Jinglin Kong, Woojong Rho, Ivan Teahulos, Arya Tavakoli, Samantha Bovaird, Shahem Merjaneh, Andrew Tan, Kiran Edwardson, Christelle Scheepers, Andy Ng, Andy Hajjar, Baly Sow, Michael Vrouvides, Andy Lee, Philippe DeCorwin-Martin, Shafqat Rasool, Jiamin Huang, Timothy Erps, Spencer Coffin, Yu Han, Srinivas Niranj Chandrasekaran, Lisa Miller, Maria Kost-Alimova, Adam Skepner, Shantanu Singh, Anne E. Carpenter, Jeffrey Munzar, David Juncker

**Affiliations:** Nomic Bio, Montreal, QC, Canada; Broad Institute of MIT and Harvard, Imaging Platform, Cambridge, MA, USA; Broad Institute of MIT and Harvard, Center for the Development of Therapeutics, Cambridge, MA, USA; Victor Phillip Dahdaleh Institute of Genomic Medicine, Montreal, QC, Canada; Biomedical Engineering Department, McGill University, Montreal, QC, Canada

## Abstract

We present the nELISA, a high-throughput, high-fidelity, and high-plex protein profiling platform. DNA oligonucleotides are used to pre-assemble antibody pairs on spectrally encoded microparticles and perform displacement-mediated detection. Spatial separation between non-cognate antibodies prevents the rise of reagent-driven cross-reactivity, while read-out is performed cost-efficiently and at high-throughput using flow cytometry. nELISA can measure both protein concentration and their post-translational modifications. We assembled an inflammatory panel of 191 targets that were multiplexed without cross-reactivity nor impact on performance vs 1-plex signals, with sensitivities as low as 0.1 pg/mL and measurements spanning 7 orders of magnitude. We then performed a large-scale inflammatory-secretome perturbation screen of peripheral blood mononuclear cells (PBMCs), with cytokines as both perturbagens and read-outs, measuring 7,392 samples and generating ∼1.4M protein data points in under a week; a significant advance in throughput compared to other highly multiplexed immunoassays. We uncovered 447 significant cytokine responses, including multiple putatively novel ones, that were conserved across donors and stimulation conditions. We validate nELISA for phenotypic screening, where its capacity to faithfully report hundreds of proteins make it a powerful tool across multiple stages of drug discovery.

## Introduction

Accurate quantification of proteins is essential for understanding the state of a biological system, as well as for translating these insights into new diagnostic, prognostic and treatment strategies.^1^ While genomics and transcriptomics offer valuable insights and have been used as proxies for protein profiling, they may not fully capture the complexity of protein dynamics due to factors such as post-transcriptional regulation, differential translation rates, protein degradation, and spatiotemporal regulation.^1^ This highlights the need for protein profiling methods that can provide a truer representation, and more direct observation, of biology. However, the advent of proteomic tools has lagged due to unique challenges with proteins, such as the large dynamic range at which they are expressed, the existence of multiple proteoforms, post-translational modifications and protein complexes, and the lack of sample amplification afforded by PCR. As a result, scientists continue to lack broadly applicable protein analysis tools that can operate at high-throughput and multiplex at large scale without compromising fidelity, precision, and, perhaps most importantly, that are affordable and accessible.

Significant efforts have been dedicated to scaling of throughput and multiplexing in proteomics.^1^ Mass spectrometry (MS)-based proteomics can be useful for discovery in many contexts, providing quantitative information for thousands of proteins and their post-translationally modified forms. However, MS has so far failed to overcome severe trade-offs between throughput and sensitivity, limiting its throughput and remaining biased towards proteins in high abundance.^1^ The sandwich immunoassay, commonly known as the ELISA (enzyme linked immunosorbent assay), can be performed in high-throughput and remains a ubiquitous tool for protein quantification from bench to clinic. Sandwich ELISAs are of special interest, being quantitative and highly specific due to the use of a capture antibody (cAb) and detection antibody (dAb) to bind the analyte and generate a measurable signal. However, multiplexing ELISA systems to capture significant portions of the proteome has been limited by reagent-driven cross-reactivity (rCR), which severely jeopardizes assay fidelity, even at low-to mid-plex levels.^2^ In fact, rCR is the major barrier to multiplexing immunoassays beyond ∼25-plex, resulting in many assay kits limiting content to ∼10-plex. rCR is caused by the mixing of non-cognate antibodies, which are combined in solution and incubated together for target detection.^2,3^ This mixing enables combinatorial interactions between all antibodies and proteins, and allows the formation of mismatched sandwich complexes, which can be formed as a result of a single non-specific binding event. These non-specific interactions increase exponentially as the number of antibody pairs in solution multiply, increasing the background noise and thereby eroding assay sensitivity. This problem has limited the multiplexing of immunoassays to <50-plex, even with intensive selection of antibodies to minimize rCR.^1,2,4^

Efforts to minimize rCR has resulted in commercially available platforms such as Olink’s proximity extension assay (PEA) technology,^5^ which does not prevent rCR but selectively reports bi-specific interactions using proximity-based DNA amplification of oligonucleotide pairs that are analyzed by sequencing, and Somalogic’s SomaScan,^6^ using Somamers (a form of aptamer) where non-specific binding is minimized through multiple capture-release steps and detection by hybridization of Somamers to DNA microarrays. Both methods allow thousands of proteins to be measured in every assay, and are increasingly used, but the conversion of protein binding into oligo products, and their analysis, is costly ($200-700 per sample), low throughput, not suited for protein target customization on-the-fly, and not designed to measure post-translational modifications or protein complexes. We previously reported methods to spatially array and separate miniature sandwich immunoassays so as to prevent antibodies from mixing and thus alleviating potential cross-reactivity.^3,7,8^ However, lengthy spotting protocols and technical challenges increased cost and limited the reproducibility and throughput, resulting in similar limitations to PEA and SomaScan. Thus, the need remained for a platform that achieves specificity at high-multiplexing without compromising other parameters such as throughput, cost-efficiency, and versatility.

Here, we introduce the nELISA, a next-generation bead-based assay platform that combines two technologies to achieve high-fidelity at large scale multiplexing. The first technology is CLAMP (colocalized-by-linkage assays on microparticles), which we introduce here for the first time. CLAMP is a novel sandwich assay design that completely prevents rCR by pre-immobilizing antibody pairs on the surface of microparticles and detects target proteins using a novel detection-by-displacement approach, both of which are enabled using DNA oligos. We showcase the versatility of CLAMP by both quantifying protein concentration and measuring post-translational modification, and protein complexes. We combine CLAMP with a large-scale encoding and decoding approach to bead-based flow cytometry assays that we previously described,^9^ creating the nELISA. As a first demonstration of the nELISA, we built a 191-plex inflammation-targeted panel that targets low-abundance cytokines, chemokines, and growth factors. We characterized the sensitivity, specificity and reproducibility of the platform, and illustrate its potential for high-throughput screening (HTS) by profiling 191 proteins in 7,392 samples in <1 week. We illustrate the ease with which the nELISA can be integrated to existing high-throughput screening (HTS) workflows by combining it with Cell Painting for phenotypic screening of a reference compound set. Finally, we demonstrate that the nELISA recapitulates hundreds of expected immune phenotypes in a single experiment, while also revealing unexpected insights with direct implications for drug discovery and development.

## Results

### CLAMP: a miniaturized sandwich assay on microparticles that avoids rCR

To overcome the rCR and throughput limitations of multiplexed immunoassays, we developed the CLAMP, which miniaturized the ELISA assay at the surface of a bead (Fig. 1). The CLAMP departs from the classical sandwich ELISA in three key ways: 1) pre-assembly of antibody pairs; 2) releasable detection antibodies; 3) conditional signal generation. First, detection antibodies are pre-assembled on their respective capture antibody-coated beads using flexible and releasable DNA oligo tethers (Fig. 1a, Step 1). Spatial separation between non-cognate antibody pairs, each immobilized on different beads, precludes non-cognate interactions. When samples are mixed with the beads, target proteins can be recognized simultaneously by the antibodies, forming a ternary sandwich complex (Fig. 1a, Step 2). Second, the releasable tethering of the detection antibodies allows for their displacement by toehold-mediated strand displacement.^10^ CLAMP uses a novel mechanism of signal transduction using a detection-by-displacement protocol based on toe-hold mediated strand displacement (Fig. 1a, Step 3). Third, detection of the fluorescently labeled displacement oligo results in conditional signal generation, as only target-bound sandwich complexes are both labeled by the fluorescent displacement oligo and remain bound at the surface of the bead; release efficiency of oligo displacement is >98% (Supp. Fig. 2). In contrast, non-displacement is not associated with a fluorescent signal, because target absence, or non-specific interaction to one of the immobilized antibodies, result in washing away of the fluorescent probe and signal, thus ensuring low background (Fig. 1a, Step 4).

**Figure 1.**
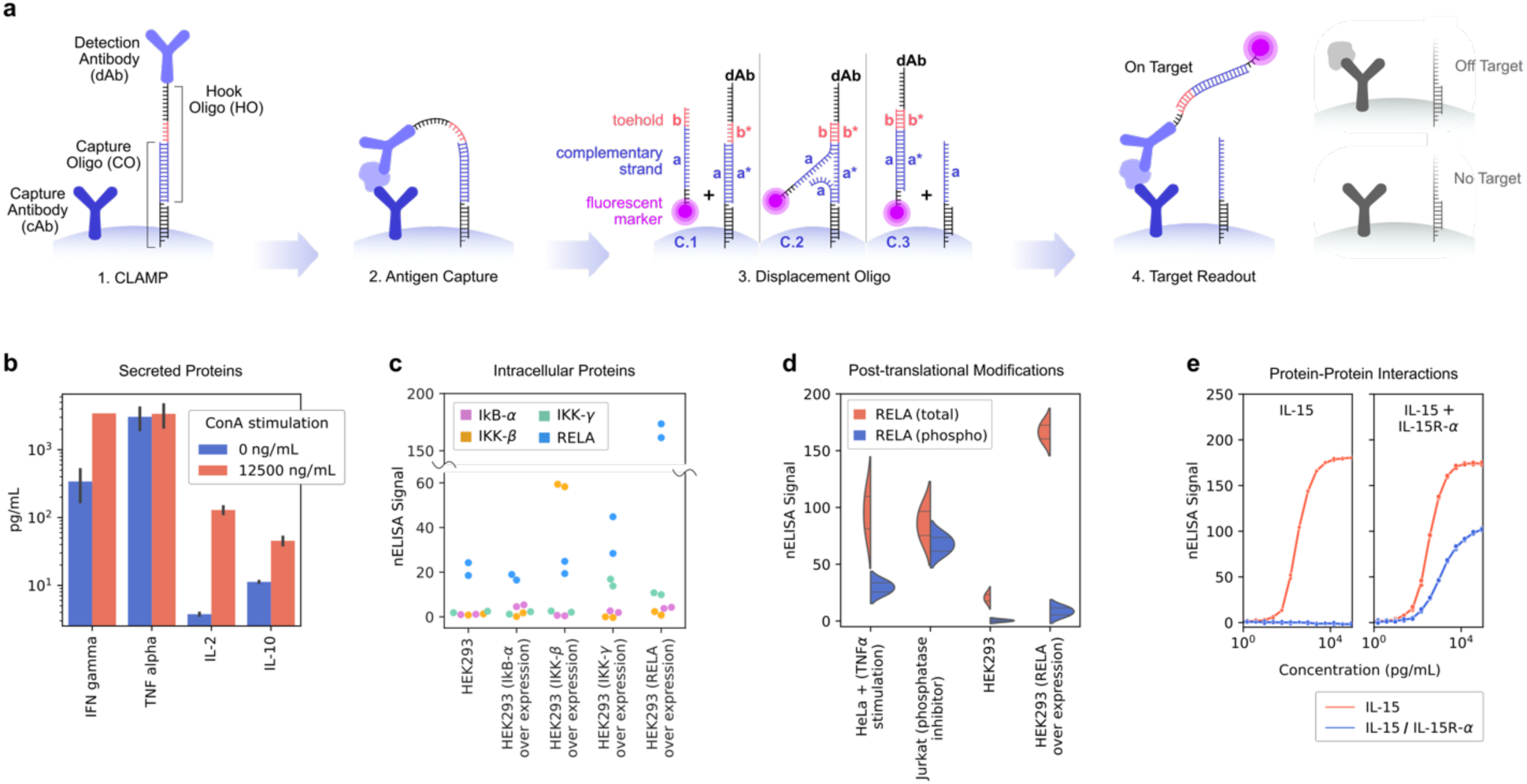
Colocalized Assay on Micro-Particles (CLAMPs). (a) Each barcoded micro-particle has pre-assembled, miniaturized ELISAs for a single target. (1) **CLAMP**: The detection antibody (dAb) is bound to a hook oligo (HO) that is tethered to the surface via partial hybridization with a capture oligo (CO) strand. (2) **Antigen Capture**: The assay is carried out by incubating the biological sample with barcoded CLAMPs generating sandwich binding in the presence of the target analyte only. (3) **Displacement Oligo**: After washing, a fluorescently labeled displacement oligo (DO) is added to displace HOs from their pre-hybridized COs via toehold mediated displacement. (4) **Target Readout**: Labeling of the sandwich complexes that remain on the surface due to target presence and sandwich formation (left, in color) or washing away of the dAb-DO hybrid in the case of off-target binding (top right) or absence of target (bottom right). (b-e) Examples of nELISA configurations enabling multiplexed measurements. (b) **Secreted Proteins**: Increased levels of inflammatory markers IL-2 and IL-10 upon ConA stimulation of PBMCs (*n*=46); error bars show standard deviation. (c) **Intracellular Proteins:** configuration is identical to that of secreted proteins, but with antibodies targeting intracellular proteins; swarm plot with nELISA measurements (*n*=2) of total levels of indicated proteins in HEK293 cells or HEK293 cells over-expressing one of the target proteins. (d) **Post-Translational Modification (PTM):** the detection antibody is specific to a PTM (for example phosphorylation); shown are violin plots with measurements (*n*=2, indicated with line) for total RELA (red) and phosphorylated RELA (blue) in cell lines with indicated treatments resulting in distinct levels of RELA expression and phosphorylation. (e) **Protein-Protein Interactions (PPI):** each antibody of an ELISA pair binds to a distinct member of a PPI; shown are 4 parameter logistic curve fits of total IL-15 (red line) and the IL-15:IL-15Ra dimer (blue line) with their respective sensors, in the presence of recombinant IL-15 (left) or IL-15:IL-15Ra (right).

During the displacement step, detection antibodies released from the beads remain at femtomolar concentrations in solution (Supp. Calculation 1), which is orders of magnitude lower than in a typical sandwich assay. This minimizes off-target complex formation. The local antibody concentration at the bead surface is high to ensure sandwich assay formation. However, the total amount of antibodies released into solution remains extremely low due to (1) the limited number of target-specific microparticles (∼50 beads per assay) and (2) the restricted antibody load per bead. As a result, detection antibody concentrations in solution are too low to drive off-target binding, preventing detectable non-specific interactions (Supp. Calculation 1).

We present proof-of-concept experiments illustrating the broad application of CLAMP for secreted and intracellular proteins, protein modification, and protein-protein interactions. CLAMP was used to detect (i) secreted inflammatory biomarkers by PBMCs upon stimulation with Concanavalin A (ConA), a well-known T-cell stimulus (Fig 1b). As with secreted proteins, (ii) intracellular targets can be detected in cell lysates using antibodies targeting intracellular proteins, as shown with HEK293 cells overexpressing intracellular IkB-α, IKK-γ, IKK-β and RELA (Fig 1c). Of note, our results confirm the well-established stabilization of NF-κB pathway components through binding partner overexpression: overexpressing either RELA or IKK-γ stabilized the other (Fig. 1c). (iii) Post-translational modifications were detected using phospho-specific detection antibodies. As expected, high levels of phospho-RELA were measured in conditions of NF-KB pathway activation due to TNFα a stimulation or treatment with phosphatase inhibitors, while phospho-RELA remained in HEK293 cells with regular and supraphysiological levels of RELA (Fig 1d). Similarly, CLAMPs distinguished between total and phosphorylated forms of IkB-α and IKK-β, with total levels increasing in response to overexpression of the protein (or its binding partners in the case of IkB-α), and higher phosphorylated levels were detected in the presence of TNFα stimulation or phosphatase inhibition (Supp. Fig. 4) (iv) Protein-protein interactions leading to the formation of protein complexes such as IL-15 and the soluble form of its cell surface receptor IL-15Ra were selectively measured using a cAb and dAb antibody for each protein in the complex, respectively (Fig 1e). Likewise, protein complexes for IL-12 p70 and IL-23 (each containing the IL-12 p40 subunit) can be detected with high specificity with CLAMPs for their respective heterodimer antigens (Supp. Fig 5). These data demonstrate the versatility of CLAMP for measuring soluble proteins, protein modifications, protein interactions in a way that captures the complex interplay between intermediates in intracellular signaling networks, as well as the secreted proteins enabling intercellular communication.

### nELISA combines CLAMP with emFRET to achieve high-throughput, large scale multiplexing

To fully leverage the multiplexing capacity enabled by the rCR-free nature of the CLAMP, we combined it with our previously-developed fluorescent barcoding technique, ensemble multicolour FRET (emFRET).^9^ The emFRET technique is designed for multiplexed bead barcoding, which enables the simultaneous measurement of multiple biological targets within a single sample. Using varying ratios of four standard spectral dyes (AlexaFluor 488 (N1), Cy3 (N2), Cy5 (N3) and Cy5.5 (N4)), thousands of unique spectral barcodes can be generated. The emFRET model compensates for Förster resonance energy transfer (FRET) occurring between spectral dyes, enabling precise prediction of the barcode universe when compared to the theoretical position (Fig 2b).

**Figure 2.**
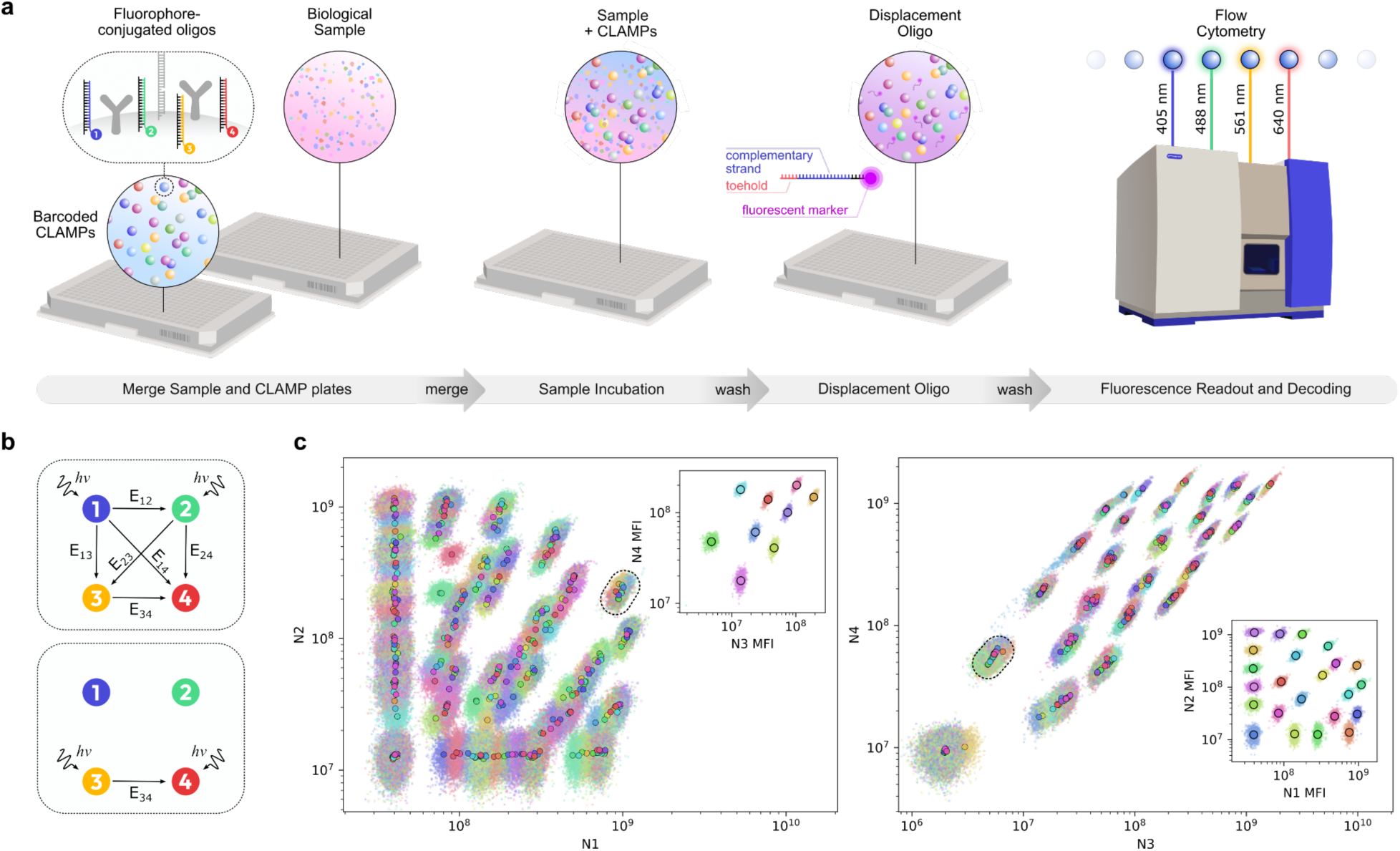
High-throughput, large scale multiplexing with nELISA by combining barcoded CLAMPs with emFRET. (a) CLAMP microparticles contain a ratio of fluorophore-conjugated oligos bound to the surface. The ratio of N1, N2, N3, N4 dyes provides a unique spectral barcode assigned to a CLAMP for a specific protein target. Following sample incubation, the addition of a displacement oligo adds a fifth dye which is used for quantification. Barcoded CLAMPs are read out by high-throughput, 4-laser flow cytometry. (b) The emFRET model compensates for inter-dye energy transfer upon excitation of each spectral dye, such as energy transfer from dye 1 to 2 (E12) upon excitation of dye 1. This enables accurate modeling of barcode positioning and decoding of each barcode to its target assay. (c) Cytometry data from a 384 spectral barcode universe. Each decoded barcode is labeled with a color. Markers (circle) of a matching color indicate the emFRET model prediction of the barcode position. Barcodes with spectral overlap in N1/N2 (left) are spectrally distinct in N3/N4 (inset shows barcodes outlined with dotted line). Likewise, barcodes with overlap in N3/N4 (right) are distinct in N1/N2 (inset).

To achieve large-scale encoding of nELISA beads, spectral dyes N1-N4 are conjugated to a DNA oligo and combined in varying ratios (from 0-70 parts per dye) using automated liquid handling to achieve a unique spectral barcode. Microparticles coated with a complementary oligo are incubated with the spectral dye mixture, which binds to the surface of the bead (Fig 2a, left). These barcoded microparticles are each assigned to a specific protein target and combined with specific CLAMP reagents for this target. Assembled CLAMPs are then pooled and dispensed into 384-well plates.

Using four fluorescent dyes with partially overlapping spectra at various concentrations, we generated 384 distinguishable barcodes (Fig 2c) for our experiments. Barcodes with overlap in N1/N2 channels are distinct in N3/N4 and vice versa. Furthermore, the fluorescence-based readout enables the use of high-throughput flow cytometry, and the use of 384-well plates to achieve high-throughput protein profiling. We developed workflows enabling 1,536 wells to be analyzed daily on a single cytometer, the highest throughput of any highly multiplexed immunoassay achieved to date, to the best of our knowledge. The number of barcodes could be readily increased to several thousand by optimizing spectral distribution and including additional dyes, so that nELISA could profile thousands of proteins without compromise on its very high throughput.

### Fidelity of assay performance is maintained at high plex

To demonstrate high-fidelity multiplexing, we assembled an inflammation-focused panel composed of 191 targets including cytokines, chemokines, and growth factors (Suppl. Table 1). We chose secreted proteins as the proof-of-concept set of proteins because they are simultaneously important to study, difficult to measure due to low abundance, and exhibit pleiotropic effects that depend on context and co-stimulations, therefore requiring broad profiling to better understand their roles ^11^. To determine whether multiplexing had any negative effect on protein quantification, we ran a head-to-head measurement comparing signals generated in single-plex vs 191-plex. For individual proteins, calibration curves were indistinguishable; across the 191-plex panel, measurements correlated with an R^2^ of 0.988 (Fig. 3b), showing that nELISA measurements are insensitive to multiplexing.

**Figure 3:**
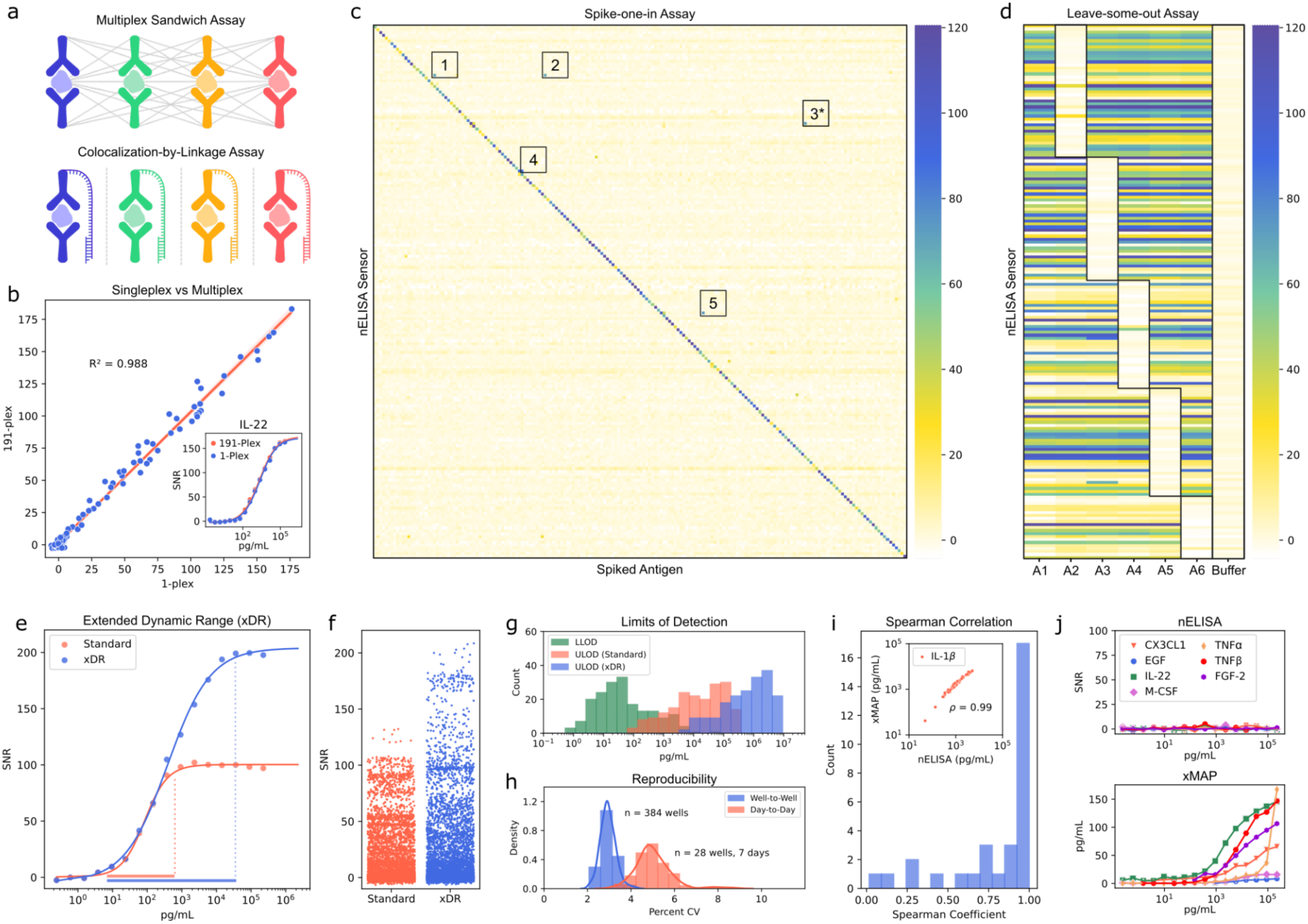
Characterization of nELISA specificity, sensitivity, and reproducibility. (a) Schematic representation of rCR in traditional multiplexed ELISAs, and its abrogation by CLAMP. (b) Correlation of SNR values for CLAMPs in 1-plex, or 191-plex format; an example standard curve is also shown (inset). (c) Spike-one-in assay: heatmap displays SNR values for each nELISA sensor in the 191-plex (y-axis) for each individually spiked antigen (x-axis); diagonal displays specific signals, cross-reactive events are numbered. (d) Leave-some-out assay: 6 pools of recombinant proteins consisting of either all the targets in the 191-plex panel (A1) or lacking a subset of targets (A2-A5) were profiled; outlined boxes indicate absence of target proteins; colours represent SNR. (e) Overlaid standard curves for 1 example CLAMP using (red) standard and (blue) extended dynamic range (xDR) protocols. (f) Distribution of SNR values for all 191-plex across 80 cell culture supernatants from stimulated PBMCs quantified using (red) standard or (blue) xDR protocols. (g) Distribution of the lower limits of detection (LLOD, green) and upper limits of detection (ULOD, red or blue) of 191 sensors using (red) standard or (blue) xDR protocols. (h) Distribution of coefficients of variation (%CV) of all nELISA sensors in repeat measurements of a single sample across wells in a single plate (well-to-well, blue) and across plates profiled on different days (day-to-day, red). (i) Cell culture supernatants from stimulated PBMCs were analyzed with nELISA and xMAP platforms; shown is the distribution of Spearman correlation coefficients for shared sensors with detectable protein concentrations, and an example of correlating IL-1β concentrations (inset). (j) Cross-reactivity comparison on xMAP and nELISA platforms: 100 recombinant antigens were pooled and spiked in cell culture media at increasing concentrations; shown are the quantification of 7 proteins which are not detectible in the sample using the nELISA (top) and xMAP (bottom).

We then screened for rCR by running ‘spike-one-in’ testing, where each recombinant antigen was separately spiked to detect any cross-reactive detection by non-cognate CLAMPs (Fig. 3c). Of the 36,000+ possible cross-reactivities (where protein x is detected when protein y is spiked), we only detected 5; of these 3 were explained by shared epitopes. For example, it was expected that an IL-12 p40-specific CLAMP would detect IL-12 p70 and IL-23, as these heterodimers both contain the IL-12 p40 subunit. It was also expected that CXCL12α and CXCL12β CLAMPs would detect both CXCL12 isoforms, as they differ by only 4 residues. Unexpectedly, the CCL13 CLAMP detected CCL17, and the CCL3 CLAMP detected PCSK9. These cross-reactivities are presumed to be due to sample cross-reactivity (as opposed to rCR), whereby both antibodies detect the same off-target and would yield cross-reactive results even in a single-plex ELISA and can be addressed by selecting alternative antibody pairs. Of note, detecting such cross-reactive events is a challenging and resource-intensive step for ELISA developers; our results suggest that the nELISA could be leveraged to accelerate and improve this process. We further validated nELISA specificity in more complex protein samples by running ‘leave-some-out’ testing, in which antigen pools lacking a subset of nELISA targets were profiled to ensure rCR remained low even in the presence of other abundant proteins in solution (Fig. 3d). These results establish the specificity and accurate quantification of our 191-plex secretome panel, demonstrating that the nELISA fidelity is unaffected by multiplexing, well beyond the traditional limit imposed by rCR.

### Characterizing dynamic range, sensitivity, and precision of nELISA

To further characterize the performance of the nELISA in 191-plex, we analyzed standard curves generated for each nELISA target. nELISA yielded sigmoidal dose-response curves similar to classical ELISA and were robust, up to 100µg/mL tested here, against the Hook effect that can affect quantification at high target concentrations in some immunoassay formats ^12^ (Fig 3e). These characteristics enabled us to extend the dynamic range (xDR) of the assay by merging concentrations obtained from profiling proteins at 2 dilutions (see Methods), without impacting the assay throughput (Fig. 3e-f). Thus, high abundance proteins with concentrations up to 10 µg/mL could be quantified accurately. At the lower end of the detection spectrum, we analyzed the sensitivity of the nELISA and found limits of detection as low as 0.1 pg/mL (Fig. 3g). The assay binding range was generally governed by antibody affinities, as we observed concordance between CLAMP and classical ELISA formats. Combined with our xDR method, the nELISA thus yielded a collective protein quantification range of 7 orders of magnitude, with each CLAMP possessing a dynamic range of 3-5 orders of magnitude.

Importantly, nELISA signals are reproducible. By measuring the same sample across wells, plates and profiling days, we calculated that the coefficient of variation (CV) of each CLAMP was ∼3% well-to-well, and 5% plate-to-plate and day-to-day (Fig. 3h). As a result of variations much smaller than that of biological assays, the nELISA is inherently repeatable, enabling large scale cell-based screening. Furthermore, the nELISA includes several layers of controls that ensure signal stability over time (Suppl. Fig. 1). Non-human targeting CLAMPs are added to every well as a Displacement Control, ensuring that CLAMPs do not yield signals in the absence of their target. In addition, Signal Normalization Controls are added, consisting of CLAMPs lacking a detection antibody, and able to directly hybridize with the Detection Oligo. We also included controls on a separate calibration plate, consisting of standard curves of every CLAMP, as well as reference samples used for quality control. The performance of these controls is consistent over time, as shown by the tight overlap of standard curves performed on different days (Suppl. Fig. 6), as well as the reproducible quantification of targets in reference samples across 42 days of nELISA runs, using 4 different lots of CLAMPs (Suppl. Fig. 7).

### Benchmarking and validation against conventional multiplex bead-based assays

To validate the ability of nELISA to quantify proteins from biological samples, we collected cell culture supernatants from peripheral blood mononuclear cells (PBMCs). We profiled them with the 191-plex nELISA, and with a 48-plex panel based on the commonly used multiplexed immunoassay platform, xMAP. Of the 36 targets shared by both platforms, 24 were detected by both platforms. Quantification of the 24 detected targets revealed that xMAP and nELISA results correlated well, with a median Spearman correlation of 0.92 (Fig 3i, Suppl. Fig. 8). Leave-some-out testing revealed that targets correlating poorly between nELISA and xMAP displayed significant rCR in the xMAP platform as off-target protein concentrations approached 1 ng/mL (Fig 3j). These results demonstrate the high specificity of nELISA in multiplexed quantification, where concentrations measured are not impacted by the level of multiplexing.

To further benchmark the nELISA against a widely used proteomic platform, we conducted a direct comparison in cell supernatant samples with Olink. Olink is a widely established proteomic platform that uses a proximity extension assay (PEA) to enable multiplexed protein quantification with high specificity and sensitivity. In our head-to-head comparison, PBMCs from four donors were treated with four concentrations of each stimulus (LPS, ConA, PolyIC, PMA/i) or left unstimulated for 24 hours. Cytokine levels were quantified using the nELISA 191-plex and the Olink Explore 384 Inflammation panel. Of the 85 targets shared between platforms, 36 were consistently detected above the limit of detection on both platforms, while 31 showed no expression on either (>95% of data points). The median correlation exceeded 95%, underscoring the strong agreement between platforms (Suppl. Fig. 9). These results highlight the robustness of the nELISA, demonstrating its ability to reproduce quantitative measurements across two of the most widely adopted proteomic technologies, xMAP and Olink.

### nELISA screening of PBMC inflammatory secretome characterizes cytokine responses

To test the utility of nELISA we applied it to characterize cytokine responses and interactions in PBMCs. These cells are a uniquely accessible cell population that can be controlled and stimulated in any number of experimental conditions. This flexibility of PBMCs can thus provide a wealth of information for diverse studies, such as functional genomics, where the tunable information provides significant advantages over snapshots available from profiling plasma/serum ^13,14^. However, the diversity of conditions that can be tested also presents a major scaling challenge, as tracking PBMC cytokine responses at the protein level has so far been limited by high cost, limited throughput and limited content of available tools. While common, attempts to track cytokine responses by transcriptomics suffer from the limited correlation between mRNA and protein levels of cytokines due to extensive post-transcriptional and post-translational regulation.

To demonstrate nELISA’s ability to overcome these barriers, we selected six different PBMC donors to capture biological variability, which we treated with four different inflammatory stimuli at multiple doses. In addition, we used a library of 80 recombinant human proteins with known immunomodulatory properties as “perturbagens” to further characterize immune responses (Fig 4a), generating 7,392 profiles of human cytokines, chemokines, and growth factors. Using UMAP as an overview of the entire nELISA dataset, distinct phenotypes clustered by stimulation condition, by PBMC donors, as well as by the concentration of stimulus used (Fig 4b). In addition, individual cytokine perturbagens with particularly strong effects created their own phenotypic clusters, including IL-4, IL-10, IL-1 RA/RN, IFNα2 and IFNβ (Fig 4b, cytokine perturbation), demonstrating the power of the nELISA for phenotypic screening.

**Figure 4:**
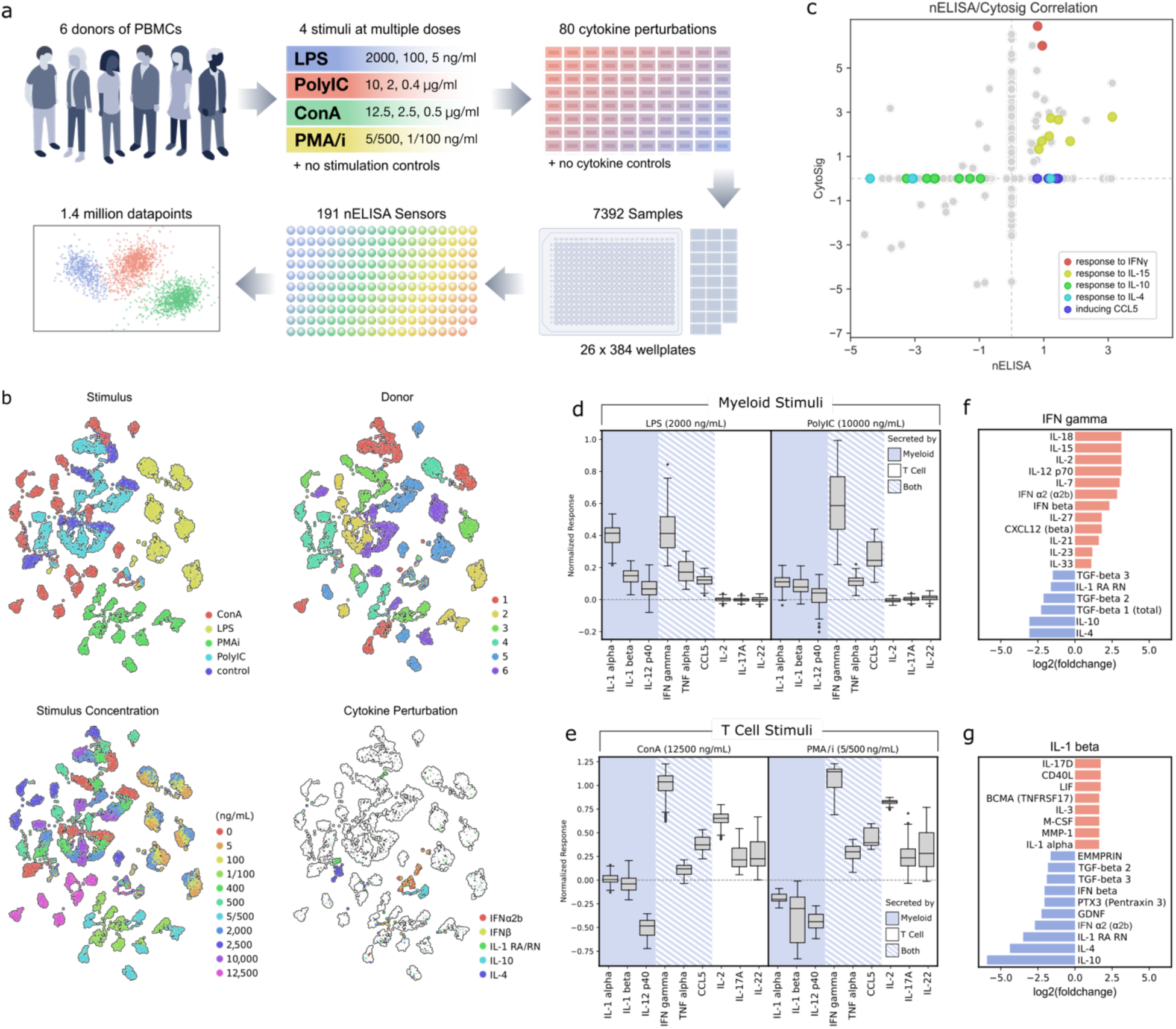
High throughput nELISA screen captures PBMC phenotypic diversity and quantifies individual cytokine interactions. (a) Screen design: PBMCs isolated from six donors were treated with inflammatory stimuli at indicated concentrations and further perturbed with 80 recombinant cytokine “perturbagens”, generating a total of 7,392 samples; after 24 hours, concentrations of 191 secreted proteins were measured in the supernatant of each sample using the nELISA. (b) UMAP dimensionality reduction of the data; data points are coloured (from left to right) by stimulation condition, by donor, by stimulation concentration, or by individual cytokine perturbagens with strong effects, as indicated. (c) Correlation between cytokine interactions detected by nELISA and CytoSig in PBMCs, according to the fold change in expression of a protein in response to a perturbagen. Examples of cytokine interactions are indicated: response to IFNγ (blue), response to IL-15 (purple), response to IL-4 (red), response to IL-10 (green), perturbagens inducing CCL5 (yellow). (d-e) PBMC response of 6 donors to myeloid and T-cell stimulus; boxplot center line shows median; limits show upper and lower quartiles; whiskers show 1.5x interquartile range; points show outliers; *n=*162. (d) PBMC expression of indicated proteins, in response to myeloid stimuli (LPS or PolyIC), in the absence of recombinant cytokine perturbagens. Normalized signals represent differences in SNR compared to control wells. (e) PBMC expression of indicated proteins, in response to T cell stimuli (ConA or PMA/i), in the absence of recombinant cytokine perturbagens. (f-g) Fold change in the expression of IFNγ (f) and IL-1β (g) in response to indicated perturbagens, across all donors and stimulation conditions.

### nELISA recapitulates classical immune responses

Using different stimulatory agents enables preferential stimulation of different subsets of cells (T cells vs myeloid cells) within PBMCs. Thus, primarily myeloid cell-derived proteins such as IL-1α and IL-1β increased in response to myeloid stimuli (Fig. 4d), whereas primarily T-cell derived proteins like IL-17A and IL-2 increased in response to T cell stimuli (Fig. 4e); proteins expressed by multiple cell types such as TNFα and IFNγ increased in response to all stimuli (Fig 4d-e) ^15^. Next, we analyzed cytokine interactions, defined as changes in expression of a given PBMC-derived cytokine in response to a given recombinant cytokine perturbagen, in any or all of the stimulation conditions, across multiple donors (see Methods). Classical pro-inflammatory cytokines such as IFNγ and IL-1β were regulated by a variety of perturbagens, in a manner consistent with the expected role of each perturbagen (Fig 4f-g). For example, IFNγ increased in response to IFNα2/β, IL2, IL-18, Il-15, and IL-7, all powerful inducers of IFNγ ^16–19^, but was suppressed by IL-4 and IL-10 ^20,21^.

To further establish that the nELISA can capture expected biology, we compared the cytokine interaction responses found with the nELISA against CytoSig, a database of consensus, data-driven, cytokine-activity transcriptomic signatures ^22^. Despite the expected differences between mRNA and protein measurements, we anticipated that well-established immune responses would be reflected in both datasets, offering an opportunity for cross-validation. We found a total of 447 cytokine interactions using our nELISA data, and 137 cytokine interactions in PBMC data from CytoSig, of which 45 were detected by both platforms. Of these, 87% agreed with respect to directionality: 29 interactions resulted in increased cytokine production, and 10 resulted in repressed responses, in both nELISA and CytoSig (Fig. 4c). These included well known responses such as (i) the induction of the IFNγ-inducible chemokines (CXCL9, CXCL10) and (ii) the potent immune-stimulatory effects of IL-15, which induced adaptive and innate immune mediators (IFNγ, TNFα, CXCL9, CXCL10, CCL5, IL-17F, IL-22, IL-1β) (Fig. 4c).

### nELISA reveals cytokine interaction insights beyond transcriptomics

Interestingly, CytoSig data lacked many hallmark effects of Th1 and Th2 modulators. For example, the potent inhibitory effect of the Th2 cytokine IL-4 on IFNγ, TNFα and IL-1β was missing in the mRNA data, as was its ability to induce CCL22 and CCL24 ^20,23–25^. In contrast, our nELISA data clearly showed these effects (8c). In fact, nELISA detected many more interactions in PBMCs than CytoSig (449 vs 137). Much of the difference stems from the inclusion of inflammatory stimuli in the nELISA dataset, enabling the detection of suppressive effects on cytokines with low baseline expression. As a result, CytoSig interactions are primarily increased expression (81%), whereas nELISA data is more balanced. This may explain why CytoSig fails to detect many of the potent anti-inflammatory effects of IL-10 on IFNγ, TNFα, IL-1β, IL-12 p40, CCL1, CCL3, CCL4, CCL5, CCL19, CXCL5, G-CSF, MMP-1, etc. ^26^ (Fig 4c).

Furthermore, CCL5, a particularly IFNγ-sensitive protein, was induced in the nELISA dataset by recombinant IFNγ and several conditions leading to increased IFNγ. However, its induction by IL-2, IL-7, IL-18, and IFNα2/β was not detected by CytoSig (Fig 4c). This may be due to post-transcriptional induction of CCL5 by IFNγ ^27,28^, or may reflect differences in experimental setup, such as temporal differences that would preclude detection of secondary effects by CytoSig. Similarly, post-transcriptional regulation likely explains differences in IL-1β regulation seen in nELISA vs CytoSig. IL-1β is primarily regulated by cleavage and release of a pre-synthesized precursor rather than de novo transcription ^29^, highlighting the need for protein-based detection to record changes. In line with this, IL-1β was the most responsive protein in the nELISA PBMC screen, responding to 35 distinct cytokine perturbagens, but only responding to IL-15 and IFNβ in the CytoSig database (Fig. 4c). This type of cytokine post-transcriptional regulation is likely the source of the disagreements between nELISA and CytoSig data. As seen in Figure 4c, there are six examples of a cytokine interaction resulting in a decrease in nELISA protein data, but an increase in CytoSig mRNA data. These interactions involve the expression levels of TNFα, IL-1α, CCL2, and CXCL1, all of which are well-studied examples of cytokines regulated at the level of translation into proteins, as well as mRNA stability, that could account for the observed differences ^30–33^. Thus, while the nELISA recapitulates well established biology captured by gene expression databases such as CytoSig, our results suggest that it may provide a more accurate representation of cell signaling states at the population level than transcriptomics.

### nELISA complements high-content phenotypic screening with mechanistic insights

Profiling the inflammatory secretome using nELISA is compatible with a wide variety of cell-based assays, by first removing the cell supernatant for nELISA and using remaining cells for any cell-based assay of interest. In fact, combining nELISA with other multiplex profiling assays probing mRNA, chromatin, or morphology can greatly increase the information gained from a single sample. We tested the nELISA in combination with Cell Painting, a high-content imaging assay that quantifies phenotypes of cells stained with six dyes that label eight cellular components, and extracts thousands of image-based features to form an image-based profile of the sample ^34^. Using samples of A549 cells (human lung adenocarcinoma) prepared by the JUMP-Cell Painting Consortium ^35^, we measured secretome profiles from the exact same samples that were then Cell Painted. These cells were treated with a library of 306 well characterized compounds from the Broad Institute’s drug repurposing library, with extensive annotations on their gene targets and mechanisms of action (MOA) ^36^.

To investigate the ability of each assay to identify compounds with shared MOA and gene targets, we first sought to leverage their existing annotations. We determined the phenotypic activity of each compound on both platforms, as measured by the ability of replicate wells of a compound treatment to match each other’s phenotypes relative to control (DMSO-treated) wells. Said differently, compounds with high self-retrieval yield a strong and distinct impact on the measured phenotype as compared to control wells. Among compounds resulting in significant phenotypic activity, nELISA and Cell Painting were able to retrieve the known MOA and gene target information (according to available compound annotations) based on shared phenotypes. MOA-retrieval rates were 21-27% and gene target retrieval rates were 15-16% on both platforms. More total compounds showed phenotypic activity by Cell Painting, which was expected due to the limited secretory capacities of A549 cells and the absence of immune stimulation (Suppl. Fig. 10). Interestingly, while some compounds’ MOAs were well predicted by both platforms, each provided better predictions for distinct subsets of compounds, resulting in retrieval of an additional 33% of MOAs by adding nELISA to Cell Painting, highlighting their complementary nature (Suppl. Fig. 10). One caveat to this analysis is that the annotations are imperfect and do not fully capture a compound’s biological effects. nELISA identified at least one case where a compound displayed a distinct phenotype from other compounds in the MOA class “CDK inhibitors”; we discovered that its unique specificity for CDK4 ^37^ yielded opposite effects on the secretome as compared to the other compounds in that MOA class (Suppl. Fig. 11). Thus, we validated our ability to run two richly informative profiling assays on the same physical samples, and to cluster individual chemical perturbagens according to their mechanism of action or gene target.

To enable exploring compounds’ relationships in an unbiased manner, without needing extensive annotations, we re-analyzed the nELISA dataset based on the similarity of the cytokine response profiles, without a priori knowledge of MOAs or gene targets (see Methods). This approach yielded additional mechanistic insights. For example, CHK inhibitors AZD7762 and 7-hydroxystaurosporine clustered closely, as did the Aurora B/C kinase inhibitor GSK1070916 (Suppl. Fig. 12). Consistent with the position of Aurora B downstream of CHK1 ^38^, the effect of GSK1070916 on A549 secretome partially recapitulated the effects of CHK inhibition, reducing the expression of IL-11, CXCL16, TNF RI, FLRG, and inducing MIF expression (Suppl. Fig. 12). Interestingly, pan-Aurora kinase inhibitors AMG900 and danusertib did not result in similar expression profiles, supporting the distinct mechanism of action of these inhibitors compared to GSK1070916 ^39^. Overall, we found many examples where nELISA data complemented Cell Painting data for MOA elucidation.

### nELISA profiles reveal cytokines with shared response profiles

To test whether the nELISA can reveal cytokines yielding similar responses independently of donor/stimulation conditions, we adapted our response profile similarity analysis pipeline to our PBMC screen. We calculated the fold change in expression of each protein in our 191-plex panel, in response to each of the cytokine perturbagens, across all donors within a stimulus condition (Suppl. Fig. 13). We also compiled effects across all stimulation conditions to capture the most reproducible perturbagen effects. UMAP clustering highlighted the most similar perturbagens (Fig 5b), with the protein expression patterns underlying these clusters shown in Fig 5a. While recombinant proteins are known to occasionally elicit an immune response due to bacterial contaminants from the manufacturing process, we observe predictable responses to recombinant cytokine perturbation, suggesting minimal if any impact from recombinant impurities.

**Figure 5:**
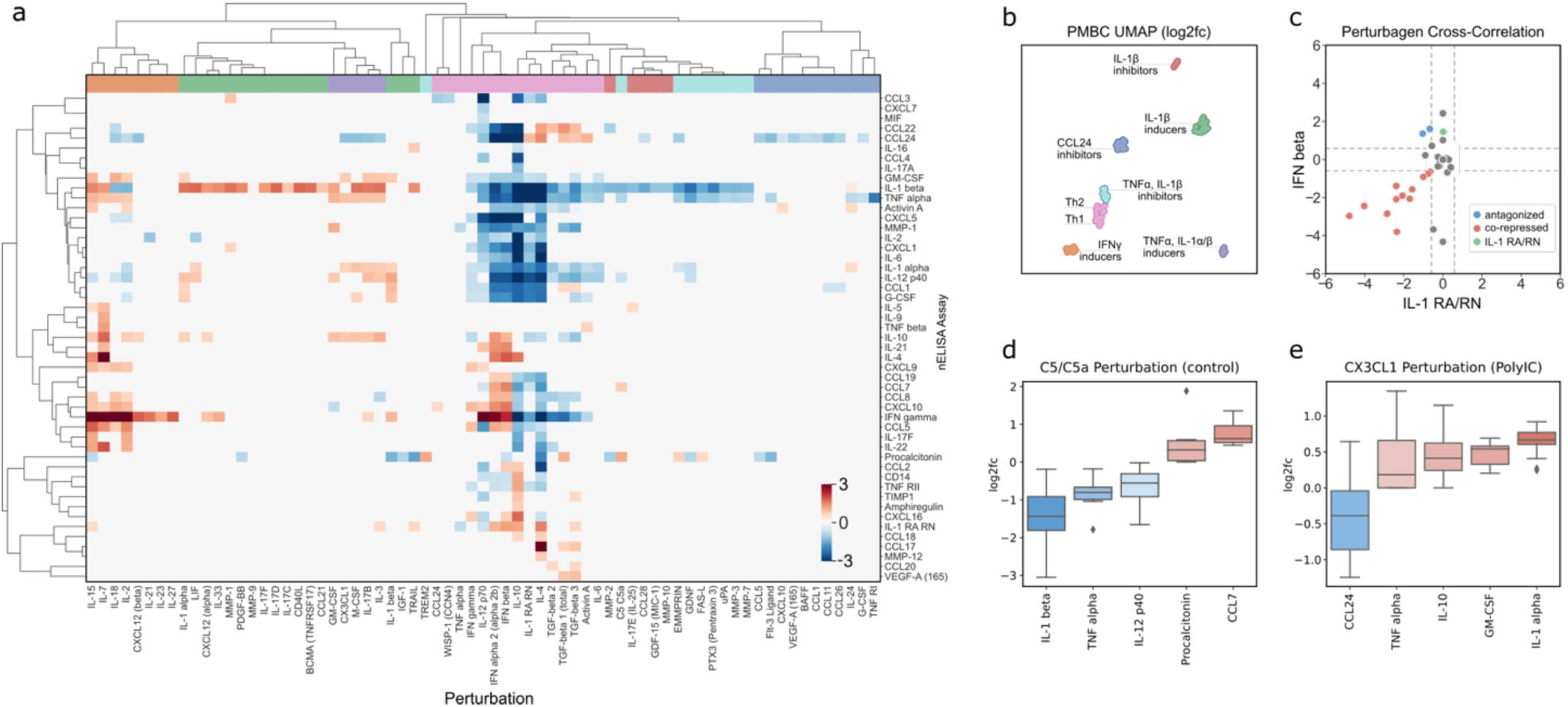
Clustering of cytokine profiles reveals phenotypically similar perturbagens and novel putative responses to recombinant chemokines. (a) Heatmap dendrogram of perturbagen effects on cytokine expression across all stimulation and donor conditions; color scale indicates fold change for each sensor over no perturbagen control. Column coloring corresponds to clusters in (b): UMAP of PBMC secretome phenotypes in response to perturbagens. UMAP of data across all stimulation and donor conditions. clusters are labeled according to shared features in PBMC secretome. (c-d) Actionable insights emerging from nELISA profiling. (c) Applicability to drug repurposing, as seen by correlating significant effects of IFNβ and IL-1 RA/RN on unstimulated PBMCs. Shown are cytokines inhibited by both perturbagens (red), cytokines (IFNγ and CXCL10) induced by IFNβ but inhibited by IL-1 RA/RN (blue), and the induction of IL-1 RA/RN by IFNβ (green). (d-e) Markers of target engagement in response to chemokine perturbagens (*n =* 6). Shown are the significantly regulated cytokines by C5a in unstimulated PBMCs (d), and by CX3CL1 in PBMCs stimulated with PolyIC (e).

We observed that the classical Th1 and Th2 immune responses clustered around the prototypical cytokines IFNγ and IL-4, characterized by induction of IFNγ/CXCL10/CCL5, and CCL17/CCL22/CCL24, respectively (Fig 5b). The proximity of these clusters is explained by the shared inhibition of innate immune responses (ex: IL-1β, TNFα, G-CSF) as seen in Fig 5b. An additional cluster that shared the suppression of IL-1β/TNFα, but had little to no effect on the expression of other cytokines, was formed by recombinant C5/C5a, EMMPRIN, GDNF, MMP-3, MMP-7, uPA, PTX3, and the soluble form of FAS-L. A distinct cluster was formed of cytokines whose main effect was to induce IFNγ expression; this included the interleukins IL-2, 7, 15, 18, 23, 27 as well as CXCL12β. Interestingly, CXCL12α clustered with another group of proteins whose main effect was to induce the secretion of IL-1β (Fig 5a). This is the largest group and included IL-1α, LIF, IL-33, PDGF-BB, sCD40L, CCL21, TRAIL and IL-17C/D/F. The remaining clusters were formed by perturbagens either inducing TNFα, IL-10 and IL-1α/β, or blocking TNFα and/or CCL24.

### nELISA uncovers cytokines with potential therapeutic implications

Interestingly, our analysis pipeline enabled us to highlight lesser-known cytokine biology, such as chemotaxis-independent effects of chemokines ^40^. Thus, CX3CL1, CCL1, CCL5, CCL11, CCL26, CXCL10, CCL24, CXCL12α/β and C5a all modulated the expression of cytokines such as IFNγ, TNFα, IL-1β, GM-CSF and IL-10, in the absence of a chemotactic gradient (Fig 5a, d). Furthermore, many of the expression changes regulated by chemokines are stimulation-dependent (Suppl. Fig. 13). These observations support a growing body of evidence for activities of chemokines beyond migration, involving signaling via distinct G-protein signals ^41^. For example, CXCL12 has been described as a co-stimulatory signal for T cells functioning through G_q_ and G_11_, rather than the G_i_ classically associated with chemotaxis ^42^. Thus, the nELISA could provide critical information in the development of therapeutic chemokines.

Notably, nELISA clustering revealed similarities between therapeutic cytokines, suggesting potential opportunities for drug repurposing. IL-1 Receptor Antagonist (IL-1 RA/RN) and IFNβ were grouped together due to their ability to inhibit innate immune responses. However, unlike IFNβ, IL-1 RA/RN did not trigger pro-inflammatory cytokines such as IFNγ and CXCL10 (Fig. 5a-c). This has implications for treating multiple sclerosis (MS), where IFNβ is commonly used and appears to work, at least partially, by increasing IL-1 RA/RN expression, despite its flu-like side effects.^43–45^ In contrast, recombinant IL-1 RA/RN (anakinra) is well tolerated and is currently being tested in a phase 1 clinical trial for MS.^46,47^ Our data showed that while IFNβ induced IL-1 RA/RN, the latter displayed only anti-inflammatory effects. Furthermore, when comparing the effects of IFNβ and IL-1 RA/RN on PBMCs, we found that IL-1 RA/RN inhibited the same cytokines and chemokines as IFNβ, except for CCL22 and CCL24, while also suppressing IFNβ-induced cytokines thought to be harmful in MS, such as IFNγ and CCL7 (Fig. 5c). These findings support the use of anakinra in MS and demonstrate the potential of nELISA protein profiling in drug discovery.

## Discussion

Although high-throughput genomic and transcriptomic methods have led to major advances in our understanding of biological processes and the molecular basis of disease, high-throughput approaches for protein profiling have lagged behind. Here, we described the development of nELISA, a protein profiling assay that leverages CLAMP to overcome the rCR limitations seen with other multiplex immunoassay systems, enabling scaling of content to 191-plex. Bead-based miniaturization of the sandwich immunoassay enabled flow cytometry-based detection, with current off-the-shelf technologies processing samples at a rate of 60 minutes per 384-well plate ^48^, resulting in a throughput of up to ∼10,000 samples per week. We demonstrate that the nELISA can scale immunoassays while maintaining or improving on their sensitivity, specificity, and dynamic range. In addition, the CLAMP technology dramatically reduces the amount of detection antibodies consumed, relative to other multiplexing formats. As reagents are the main driver of the high cost associated with protein profiling tools, nELISA assays benefit from significant advantages in cost efficiency.

Given that protein-level information is a more accurate representation of cell states and function than transcriptomics, developing improved high-throughput methods for protein profiling is of critical importance. Here, we show that nELISA can deliver insights that may not be accessible at the mRNA level. For example, the key inflammatory mediator IL-1β was the most regulated protein in our screen; because it is primarily regulated at the post-translational level, these responses were not reflected in the CytoSig database and would not be expected to be captured by transcriptomic assays. Secondary effects are also more difficult to discriminate in transcriptomic datasets, as they preclude repeat sampling to separate early events from those occurring later. In contrast, secretome profiling is compatible with repeat sampling from the same well, enabling the tracking of cellular phenotypes over time and the deconvolution of primary effects from those further downstream. Such an experiment could enable unambiguous determination of whether CCL5 is induced downstream of IFNγ, which would require an unwieldy multiplication of samples for methods requiring cell lysis or fixation, such as transcriptomics.

We applied the nELISA to HTS, where we demonstrate that it can report on the activity of small molecules and recombinant proteins. We foresee that the nELISA could be used to characterize all kinds of perturbagens and cell types, including small CRISPR gene modulation, therapeutic antibodies, bi-specific engagers, and CAR T cells. We showed that the nELISA captures expected cellular phenotypes and biological interactions that provide critical insights for drug discovery. For example, in our PBMC assay, the clear separation of responses by donor suggests powerful uses for the nELISA in functional genomics if scaled to larger cohorts, as done in the Human Functional Genomics Project ^13^.

Importantly, we showed that nELISA can be easily combined with other cell-based assays, such as Cell Painting, to provide additional biological insights. nELISA and Cell Painting provided complementary phenotypic insights, with the nELISA also providing mechanistic insight. This highlights the potential for combining nELISA with a wide range of cell-based assays, including transcriptomics, functional assays such as cell killing assays common in immuno-oncology settings, or cell surface staining experiments prevalent in immunology, to cost-effectively generate additional information from critical screens.

nELISA-based quantification of changes in the expression of secreted proteins in response to chemokines, as reported here, may have particularly useful applications in the development of therapeutic chemokines: indeed, screening with a chemotaxis assay is significantly more challenging, with lower throughput and signal-to-noise, than screening on the basis of changes in protein expression ^49,50^. Should these changes in protein expression constitute bona fide markers of target engagement, they could be leveraged for many standard drug discovery applications that suffer from the limitations of chemotaxis assays, such as large combinatorial screens, SAR studies, pharmacodynamic studies, etc. Additionally, nELISA clustering could be used to identify similarities between therapeutic interventions, such as seen in the IL-1 RA/RN and IFNβ example described above, which may inform drug repurposing efforts.

As a cytometry-based technique, nELISA provides significant cost benefits for high-throughput, high multiplex proteomics compared to mass spectrometry or approaches using sequencing-based readouts. Mass spectrometry for high-plex is costly due to extensive sample preparation, expensive reagents, and sophisticated instrumentation, which contribute to both the financial and time investment required for large-scale studies. As discussed earlier, the conversion of protein concentration to oligo concentration via somamers and PEA, and readout by DNA microarrays and sequencing, respectively, are procedurally complex, costly, and lengthy. By contrast, cytometry enables more direct and streamlined multiplexed protein detection. Its ability to simultaneously measure multiple analytes in a single sample reduces instrumentation time and reagents, significantly lowering the cost per sample and improving overall efficiency. Notably, nELISA stands out as the only high-multiplex proteomic method that uses cytometry as the readout. While conventional bead-based immunoassays use cytometry for readout, cross-reactivity between assays and panels requires profiling a sample multiple times to combine panels into higher plex. nELISA leverages the specificity of CLAMP with the ability of cytometry to rapidly measure multiple proteins in parallel, offering a more cost-effective solution for applications such as biomarker discovery and clinical diagnostics.

In conclusion, the nELISA’s high quality, uncompromising data at low cost will facilitate its adoption for high-throughput applications such as drug discovery. The self-contained nature of each CLAMP sensor, and the extensive barcoding universe accessible to the nELISA allows simple removal, exchange and addition of new sensors against new targets, and will enable the platform to expand well-beyond the 191-plex shown here, resulting in additional proteome coverage and new potential applications such as the detection of post-translational modifications, or protein-protein interactions showcased here, so long as antibodies are available. The 191-plex inflammation-focused panel described here serves to demonstrate the performance of the nELISA and paves the way for eventual profiling of the secretome, as well as intracellular proteins, protein-protein interactions, and signaling pathways in a high-throughput, cost-effective manner for cells, organs-on-a-chip, organoids, and tissues alike.

## Supporting information

Supplemental Protocol

## Acknowledgements

We thank Daniel Graham and Sarah Headland for their invaluable insights into the biology captured by nELISA profiling of PBMCs. We also thank the High-Throughput Screening Core Facility of the Institute for Research in Immunology and Cancer of the Université de Montréal for culturing and high-throughput screening of PBMCs.

## Data and Code Availability Statement

Cytokine Signaling (CytoSig) data for PBMCs is publicly available through the NIH National Cancer Institute at https://cytosig.ccr.cancer.gov/. The nELISA data and analysis code of 191-plex protein profiling in PBMCs are available in a public GitHub repository: https://github.com/nplexbio/nELISA-PBMC.

nELISA and Cell Paint data for A549 cells are available in a public GitHub repository through the Broad Institute’s Cell Paint Consortium: https://github.com/carpenter-singh-lab/2024_Kalinin_mAP/tree/v0.1.0/experiments/5_nelisa.

For inquiries regarding the A549 dataset, please contact Anne Carpenter. For questions about the PBMC dataset, please contact Milad Dagher.

## Methods

### Recombinant Sample Generation

Recombinant protein stocks and protein pools (Nomic Bio) were prepared in Sample Buffer, consisting of RPMI + 10% FBS, containing 1-191 proteins at 100 ng/mL per protein. For standard curves in singleplex and in multiplex, individual protein stocks and protein pools were serially diluted in Sample Buffer from 100 ng/mL to 0.1 pg/mL per protein. For “Spike-1-in” assays, individual protein solutions were prepared at 10 ng/mL. For “Leave-some-out” assays, protein pools containing 130-191 proteins at 100 pg/mL per protein were prepared in Sample Buffer. Each pool contained either all the targets in the 191-plex panel or lacked a subset of targets in the 191-plex panel. For cross-reactivity comparisons with xMAP, protein pool A4 was serially diluted from 200 ng/mL to 0.1 pg/mL (per protein), and aliquots were stored at -80oC for profiling by nELISA and xMAP platforms. For reproducibility testing, a reference cell culture supernatant sample was distributed across the wells of four 384-well plates; for each plate, nELISA profiling was performed on a different day, and variation was calculated across wells on the same day, and across plates on different days.

### Cell Culture

#### PBMCs

Frozen PBMCs from healthy donors (StemCell) were thawed in 37oC water bath and transferred to 50mL Falcon tube with 40mL pre-warmed medium (RPMI + 10%FBS), centrifuged 10min@200g, then resuspended in 10mL pre-warmed medium. Viability was assessed by trypan blue exclusion (>95% viability for all donors) and 50,000 viable cells (25uL at 2M cells/mL) were transferred to each well of a 384 well plate, containing 50uL per well of pre-warmed media +/-stimulus +/-perturbagens, and incubated for 24h at 37oC. Perturbagens (Nomic Bio) were present at 50 ng/mL. For stimulation conditions, LPS (InvivoGen) was present at 5 ng/mL, 100 ng/mL or 2000 ng/mL; PolyIC (InvivoGen) was present at 400 ng/mL, 2,000 ng/mL, or 10,000 ng/mL; ConA (InvivoGen) was present at 5 ng/mL, 2,500 ng/mL or 12,500 ng/mL; for PMA/i (InvivoGen), PMA was present at 1 ng/mL or 5 ng/ml, while ionomycin was present at 100 ng/mL or 500 ng/mL.

Preparation, incubation and collection of cell supernatants from PBMCs was performed at the High-Throughput Screening Core Facility of the Institute for Research in Immunology and Cancer of the Université de Montréal. After 24h, supernatants were collected, aliquots were frozen and shipped on dry ice for nELISA profiling at Nomic’s facilities or xMAP profiling at EVE Technologies facilities.

#### A549

A549 cell culture was performed at the Broad Institute’s Center for the Discovery of Therapeutics (CDoT). Cells were seeded in 4 replicate 384 well plates and cultured for 24h in DMEM + 10% FBS in the presence or absence of reference compound library. Compounds were present at a single dose (5uM). Supernatants were collected, frozen, and shipped on dry ice to Nomic’s facilities for nELISA profiling. Cells were fixed and stained for Cell Painting as previously described ^51^.

#### HEK293, HELA, and Jurkat

Cell lysates were purchased from commercial sources: HEK293 and HEK293-RELA overexpression lysates were from Origene, TNFalpha-treated HeLa extracts were from Bio-Rad, and phosphatase treated Jurkat cells were from Cell Signaling Technologies.

### DNA Oligos and Tethers

All DNA Oligos and tethers were purchased from BioIVT (Westbury, NY, USA).

#### Barcoding Oligos (BO)

BO were acquired with sequence: 5’-CACCGCCGCC ACAAAAAAAAA-[Dye]-3’ and a fluorescent dye (AlexaFluor 488, Cy3, Cy5, Cy5.5) at the 3’ end. BO are annealed to a biotinylated oligo on the surface of nELISA beads (with sequence 5’-Biotin-TTTTTTTTT GTGGCGGCG GTG-3’) in 10 μM in phosphate buffer saline (PBS)+350 mM NaCl.

#### Capture Oligo (CO)

CO have sequence 5’-Biotin-TTTTTTTTT GTGGCGGCG GTGATTGGT TATTGAGAG TTTATG-3’ and bind to the nELISA bead with a biotin terminated 5’ end.

#### Hook Oligo (HO)

HO with sequence 5’-ThioMC6-D/TTTTTTACT TTTCAACCA CCACTCAAC CATATTCAA AGCTTACGA TGCCGACTC ATTCGCCAT AAACTCTCA ATAACCAAT-3’ are terminated with a thiol modifier C6 CE-Phosphoramidite (ThioMC6) at the 5’ end. HO are conjugated to a primary amine on detection antibodies (dAb) by reduction of the HO thiol modifier with dithiothreitol (DTT) and activation with sulfosuccinimidyl 4-(N-maleimidomethyl)cyclohexane-1-carboxylate (sulfo-SMCC) to form a reactive NHS-ester. 40 μL of 30 μM thiol-modified HOs are reduced in 200 mM DTT in PBST at 37° C for one hour. Reduced oligos are (i) buffer exchanged into PBS pH 7.0 using a Zeba desalting spin-column (7K MWCO, Thermo), (ii) activated for 10 min using 8 μL of 9 mM sulfo-SMCC dissolved in 80% PBS pH7.0 and 20% anhydrous dimethyl sulfoxide , (iii) buffer exchanged again into PBS pH 7.0 to remove excess sulfo-SMCC, and (iv) a 1-10 μL fraction is reacted with 10 μL of 1 mg/mL antibodies. The reaction is left at room temperature for 1 hr and incubated overnight at 4°C. The conjugates are purified in two steps, an antibody and a DNA purification step, respectively.

### Protein Profiling

Samples of recombinant proteins or cell culture supernatants profiled using the nELISA were frozen and shipped on dry ice to Nomic’s Montreal facilities and profiled using the 191 nELISA assay targets and standard protocols. Barcoded CLAMPs were pooled and dispensed into 384-well plates at a concentration of approximately 200 beads per target per well. Sample plates were profiled by standard nELISA assay protocol (Supplementary Protocol) in batches of four plates per day. Cytometry data were spectrally decoded using emFRET to map each barcode to its target protein.

For cell lysates, protein profiling was performed by standard nELISA Assay Protocol, but with the following modifications: samples were diluted 2-fold in M-PER (Thermo Fisher Scientific) containing 1x protease/phosphatase inhibitor tablet (Thermo Fisher Scientific), 20mM EDTA, 20mM EGTA, and 100 ug/mL salmon sperm DNA.

A batch of plates containing a dilution series of calibration standards of recombinant antigens (96-wellplate format) was generated to be run with each batch of sample plates on the day of profiling. Recombinant antigens (x191) were divided into 5 groups, pooled, and diluted from 225 ng/mL in a 16-point dilution series. Media only blanks (x8) and reference samples (x8) were also included on the calibration standard plate. Calibration standards were fitted to a 4-parameter logistic curve to derive pg/mL values from fluorescence units.

Samples profiled using the xMAP platform were shipped frozen to EVE technologies (3415 A 3 Ave NW, Calgary, AB T2N 0M4) and analyzed using the Luminex Human Multiplex Cytokine Array / Chemokine Array 48-Plex (HD48) and standard protocols.

### Extended Dynamic Range Protocol

To extend nELISA dynamic ranges, samples were profiled at 2 dilutions, 2X and 50X. Standard curves were extended by stitching the linear ranges of a real standard curve and a virtual, 25x diluted, standard curve. The sum of interpolated protein concentrations from the 2X and 50X dilutions was used to derive a single protein concentration for each sample.

#### Data analysis pipeline

To identify compounds with significant effects on A549 cells, we calculated the fold change in expression of each protein in the 191-plex over the levels in control wells. Proteins with a fold change > 1.5 and p < 0.05 by Student T-test in any sample were considered significant; considering the exploratory nature of the experiment, no statistical correction was performed for multiple testing. Using the median value of the significant responses, clustergrams with hierarchical clustering were generated using cosine similarity (python package: seaborn). UMAPs of the median fold change values were generated using cosine similarity dimensions=2, spread=1.3, minimum distance=0.2, nearest neighbors=4, python package: scanpy).

To identify cytokine interactions in our PBMC assay, we accounted for stimulus- and donor-specific effects as follows. For each perturbagen in each stimulus/stimulus concentration condition, the median concentration of each secreted protein across donors was divided by the median concentration of donors in the same stimulus/stimulus concentration condition, but the absence of perturbagen, to obtain the fold change of all targets in response to all perturbagens in each condition. Significant cytokine interactions were defined by a fold change > 1.5 and p < 0.05 by Student T-test; considering the exploratory nature of the experiment, no statistical correction was performed for multiple testing. In addition, for all significant cytokine interactions, the median fold change across all stimulation conditions was calculated to identify cytokine interactions common across stimulus conditions. Clustergrams with hierarchical clustering were generated using cosine similarity and the default parameters included in the Seaborn python package. UMAPs were generated using cosine similarity with dimensions=2, spread=0.9, minimum distance=0.8, nearest neighbors=4 and the scanpy python package.

To correlate significant cytokine interactions with the CytoSig database, we identified recombinant perturbagens and responding genes/proteins that were shared in both datasets to limit comparisons to shared experimental conditions. We also limited comparisons to experiments compiled in CytoSig that were generated using PBMCs to avoid cell-type specific distinctions. Furthermore, only “high confidence” datasets were included in our analysis. For each cytokine interaction, consisting of a recombinant perturbagen and a resulting significant Fold Change in the expression of a PBMC-derived cytokine, nELISA and CytoSig results were correlated.

### Evaluating retrieval performance of nELISA and Cell Painting

We use average precision (AP) to report the ability of nELISA and Cell Painting to predict chemical mechanisms of action. Average precision is an information retrieval metric that evaluates the effectiveness of a ranking system by calculating how accurately the system ranks items based on their relevance to a query.

We use average precision for two different tasks 1) the ability of each perturbagen to retrieve its own replicates from among negative control profiles (phenotypic activity: “self-retrieval”), and 2) the ability of each perturbagen to retrieve its sister compounds (i.e., compounds that share at least one common MOA or gene target) (phenotypic consistency: “MOA retrieval” and “gene target retrieval”). We note that many compounds have multiple MOA and gene annotations rather than just one of each; in addition to the fact that annotations can be incorrect and incomplete, this makes the retrieval problem challenging and retrieval rates low for sets of compounds like ours that were not chosen for selectivity. Formally, AP is the weighted mean of precision values across all ks, where k is the number of neighbors for a given class. The definition of class varies – each *perturbation* is the class while computing AP for task 1, and each *MOA* or *gene targeted by the compound* is the class for computing AP for task 2.

We measure similarity between perturbations using cosine similarity and develop the rank-ordered similarity of all other samples to the query using this metric. Finally, we average AP per class, termed *mean Average Precision (mAP)* (for that class). For task 2, we first filter out those perturbations that cannot be retrieved relative to negative controls in task 1 and remove classes with only a single member. To set the threshold, we first calculate the p-values of each mAP in task 1 using a permutation test; compounds with a significance level of less than 0.05 are discarded. We summarize the success rate of a task by calculating the fraction of classes that have a p-value > 0.05.

### Contributions

M.D. and D.J. conceptualized the method. M.D., G.O., N.R., J.K., A.N, and J.M. developed the methodology and conceived the nELISA investigations. M.D., J.K., W.R., I.T., A.T., S.B., S.M., A.T., K.E., C.S, A.H., B.S., M.V., S.R., J.H., T.E, S.C., and J.M. performed the investigations. Y.H., S.N.C., L.M., M.K.-A., A.S, S.S., and A.E.C. conceived, developed, and performed the Cell Painting investigations and analyzed the associated data. G.O., A.L., and P.D.M. analyzed the nELISA data and developed associated software. M.D. and N.R. drafted the original manuscript. M.D., N.R., A.E.C, and D.J. reviewed and edited the manuscript. M.D. and D.J. acquired funding and supervised the study.

