## Supplemental Figures and Tables for "nELISA: A high-throughput, high-plex platform enables quantitative profiling of the inflammatory secretome"

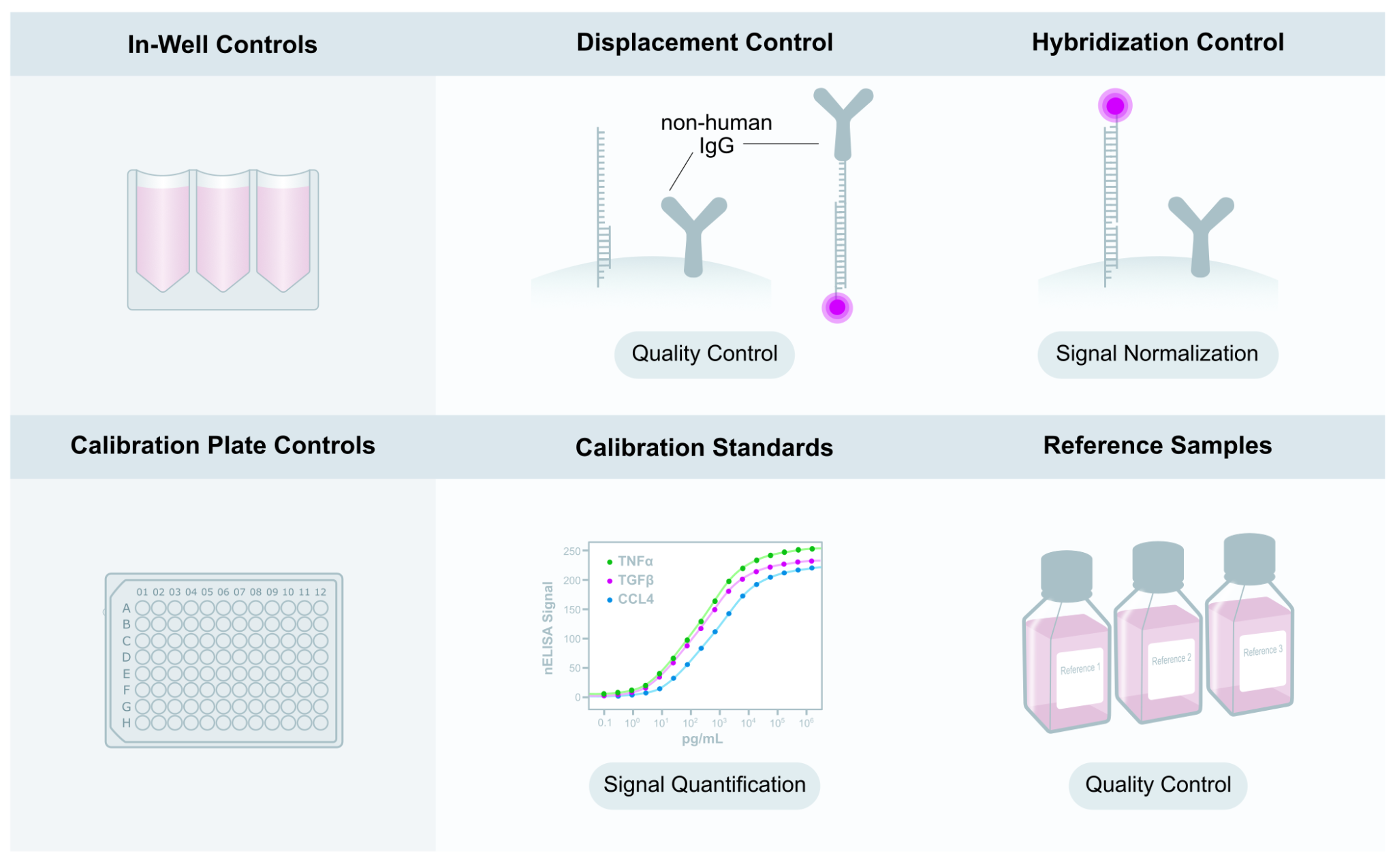

#### Suppl. Fig. 1: nELISA controls. (top) nELISA profiling includes internal controls consisting of beads with modified architectures capturing different aspects of the detection and serving as controls in each well to maintain signal consistency across wells and plates. Displacement controls consist of CLAMPs where capture and detection antibodies are specific to a non-human antigen that should not be present in any sample and are used for quality control. Signal normalization controls consist of beads with no detection antibody, directly binding to the displacer oligo. (bottom) nELISA also includes external quality controls that are run on separate plates which include calibration curves and biological reference samples.

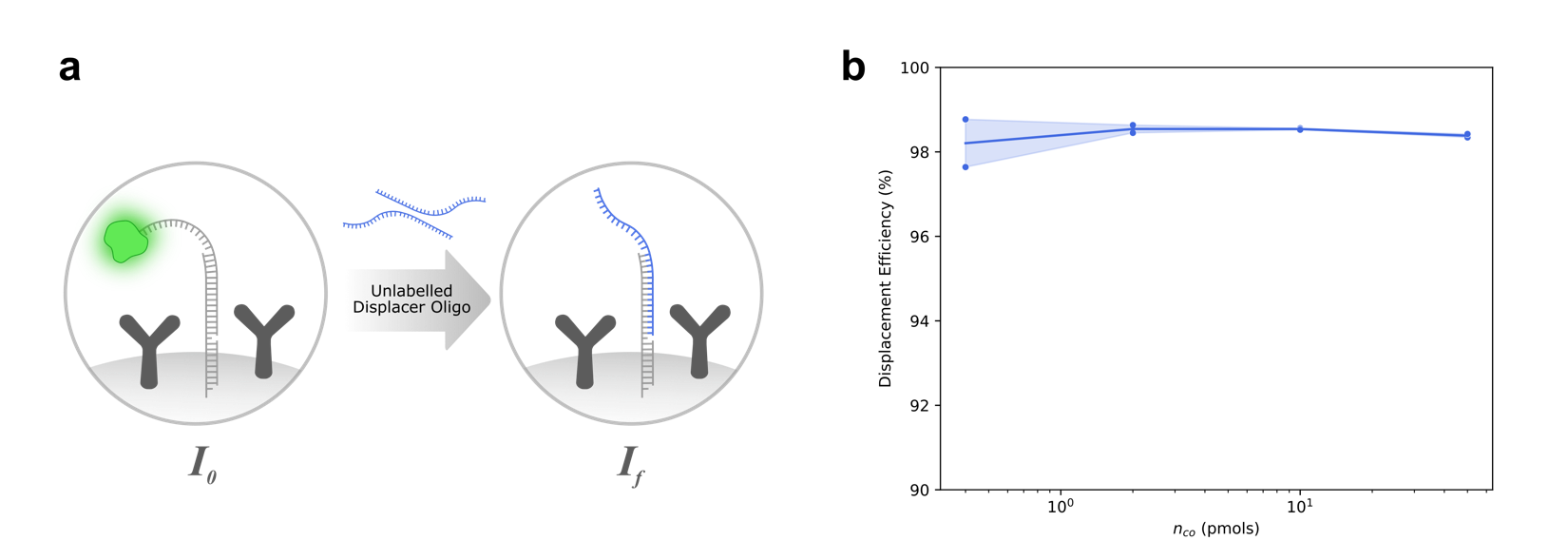

**Suppl. Fig. 2:** **Release efficiency with respect to capture oligo density.** a) release efficiency experimental design. The capture oligo on nELISA beads is annealed to a fluorescently labelled oligo. The fluorescent oligo was untethered from the bead via toehold mediated displacement with an unlabeled displacer oligo. Fluorescence of the beads before release (l_o_), after release (l_f_), and the background level (I_B_) was measure by flow cytometry (ZE5; BioRad). The release efficiency was calculated as the ratio of (l_o_ - 1_f_) to (I_o_ - 1_B_). Increased density of capture oligos had a negligible effect on release efficiency, which was consistently above 98%.

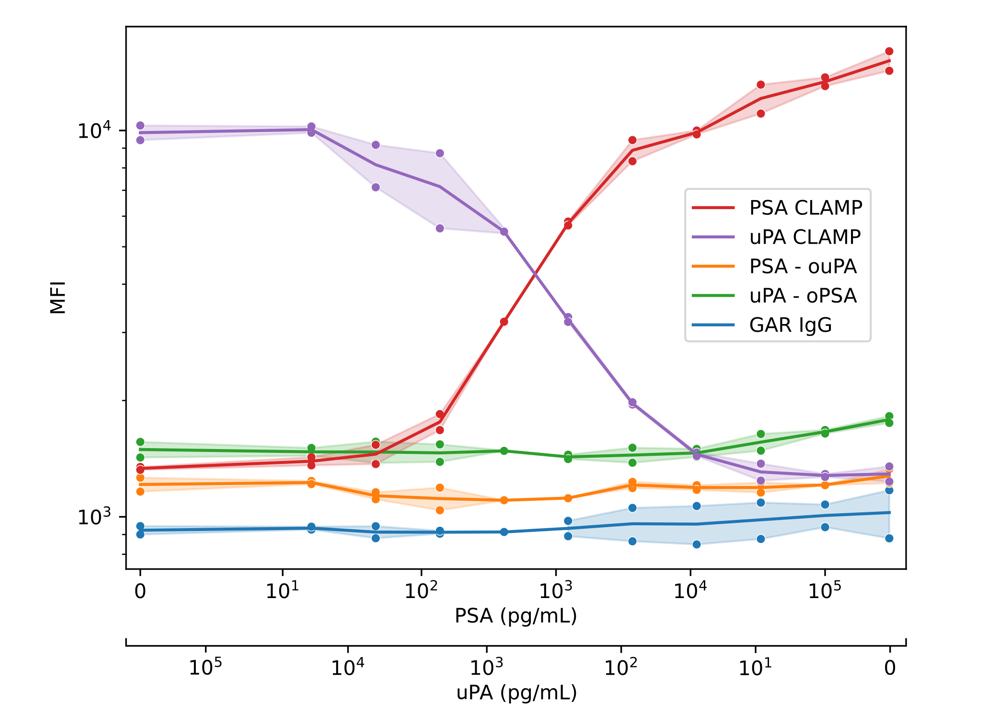

**Suppl. Fig. 3: 2-plex Cross-Reactivity Model.** The CLAMPs for PSA and uPA are compared to CLAMPs with mismatched detection antibodies, specifically the PSA capture antibody paired with the oligo-conjugated uPA detection antibody (PSA-ouPA), and vice versa (uPA-oPSA). A CLAMP using goat anti-rabbit (GAR) antibodies serves as a negative control. The total concentration of PSA and uPA is kept constant, with one antigen increasing while the other decreases. Both PSA and uPA CLAMPs demonstrate high specificity for detecting their respective antigens. In contrast, the negative control and the mismatched CLAMPs (with incorrect capture-detection antibody pairings) fail to detect either antigen in a concentration-dependent manner.

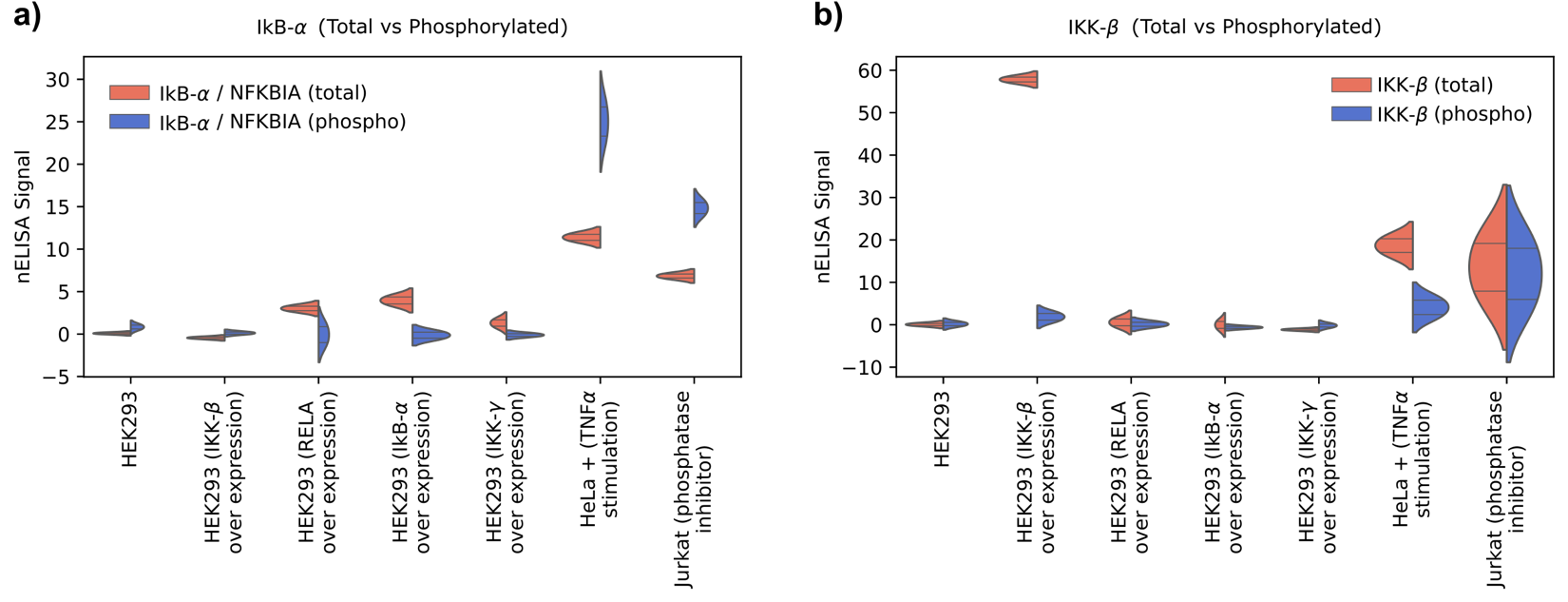

**Suppl. Fig. 4:** **Specific detection of total and phosphorylated proteins across cell types with differential expression using CLAMP sensors**. (a) Violin plots display the detection of total IκB-α (red) and phosphorylated IκB-α (blue). Total IκB-α is elevated in HEK293 cells engineered to overexpress IκB-α. As expected, phosphorylated IκB-α is elevated in Jurkat cells treated with a phosphatase inhibitor, which prevents the dephosphorylation of IκB-α, leading to its accumulation. Phosphorylated IκB-α is also elevated in HeLa cells stimulated with TNF-α, a potent activator of the NF-κB signaling pathway, which induces rapid phosphorylation as part of the pathway’s activation cascade. (b) Violin plots display total IKK-β (red) and phosphorylated IKK-β (blue) levels across the same cell types. Total IKK-β is elevated in HEK293 cells engineered to overexpress IKK-β, whereas phosphorylated IKK-β is elevated in TNF-α-stimulated HeLa cells, and phosphatase inhibitor-treated Jurkat cells.

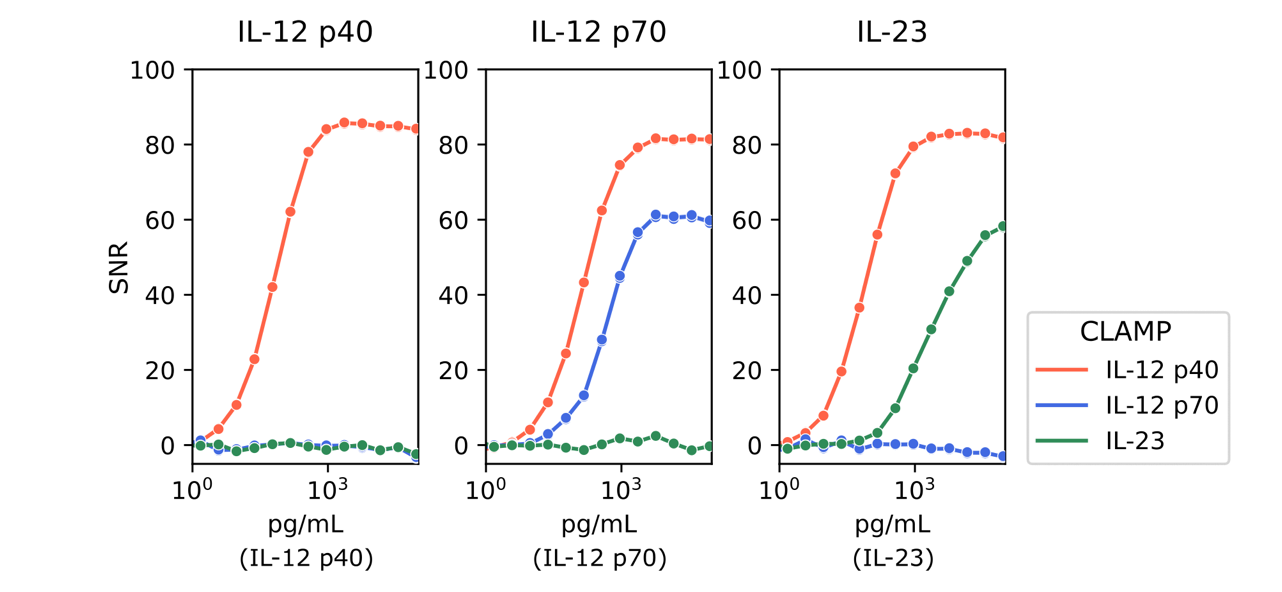

**Suppl. Fig. 5: Protein-Protein Interactions.** Example CLAMPs for detecting protein-protein interactions (PPI) with antigen dilution series for IL-12 p40, IL-12 p70, and IL-23 (left to right; plot title indicates antigen present). Both IL-12 p70 and IL-23 share the IL-12 p40 subunit. IL-12 p70 is a heterodimer composed of IL-12 p35 (IL12A) and IL-12 p40 (IL12B), while IL-23 is a heterodimer composed of IL-12 p19 (IL23A) and IL-12 p40 (IL12B). As expected, the IL-12 p40 CLAMP detects the IL-12 p40 antigen alone and also identifies the IL-12 p40 subunit within IL-12 p70 and IL-23 complexes. Conversely, the IL-12 p70 and IL-23 CLAMPs exhibit high specificity, detecting only their respective heterodimer antigens. These results highlight the capability of the CLAMP technology to precisely detect protein-protein interactions with subunit-level specificity.

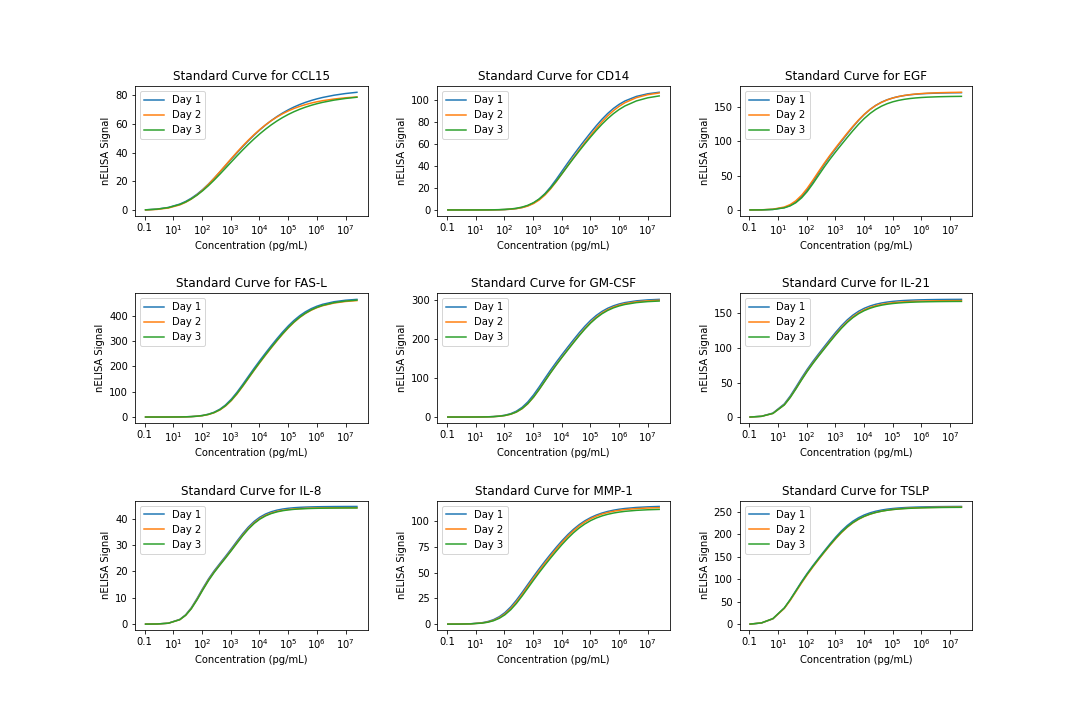

**Suppl. Fig. 6:** **Example of calibration curves.** Calibration curve fits for nine CLAMP targets shown across 3 days validation the reproducibility and consistency of nELISA assays.

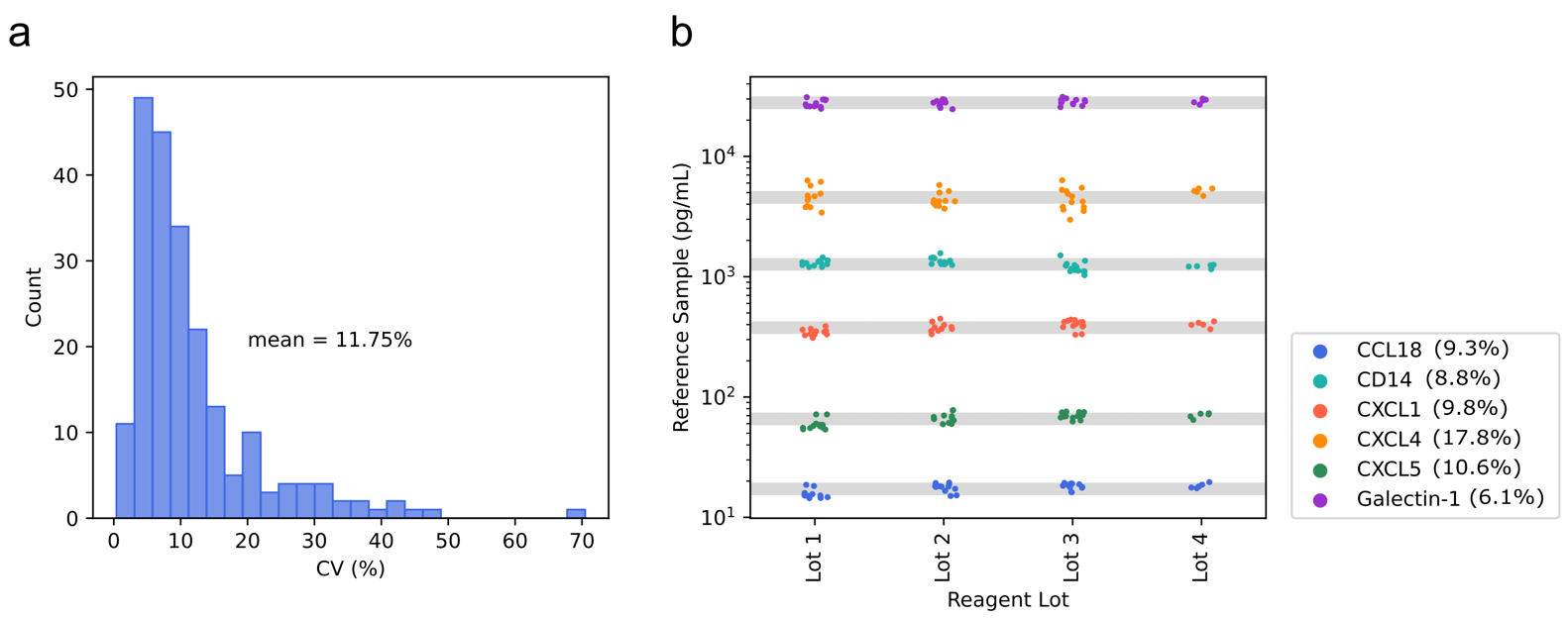

#### Suppl. Fig. 7: Measurement consistency of nELISA reference samples over time and across lots. Four lots of nELISA beads were used to repeatedly profile a reference sample included in every nELISA run. A) Histogram of CV measurements across all analytes detected in reference samples. The mean CV across analytes is 11.75%, with a median CV of 8.7%. b) Shown are measured protein concentrations for a subset of indicated analytes over 42 different days of profiling; shaded area represents 20% variation (+/- 10% from average measurement), indicating minimal variation between lots and consistency over time. Legend shows the analyte and corresponding CV.

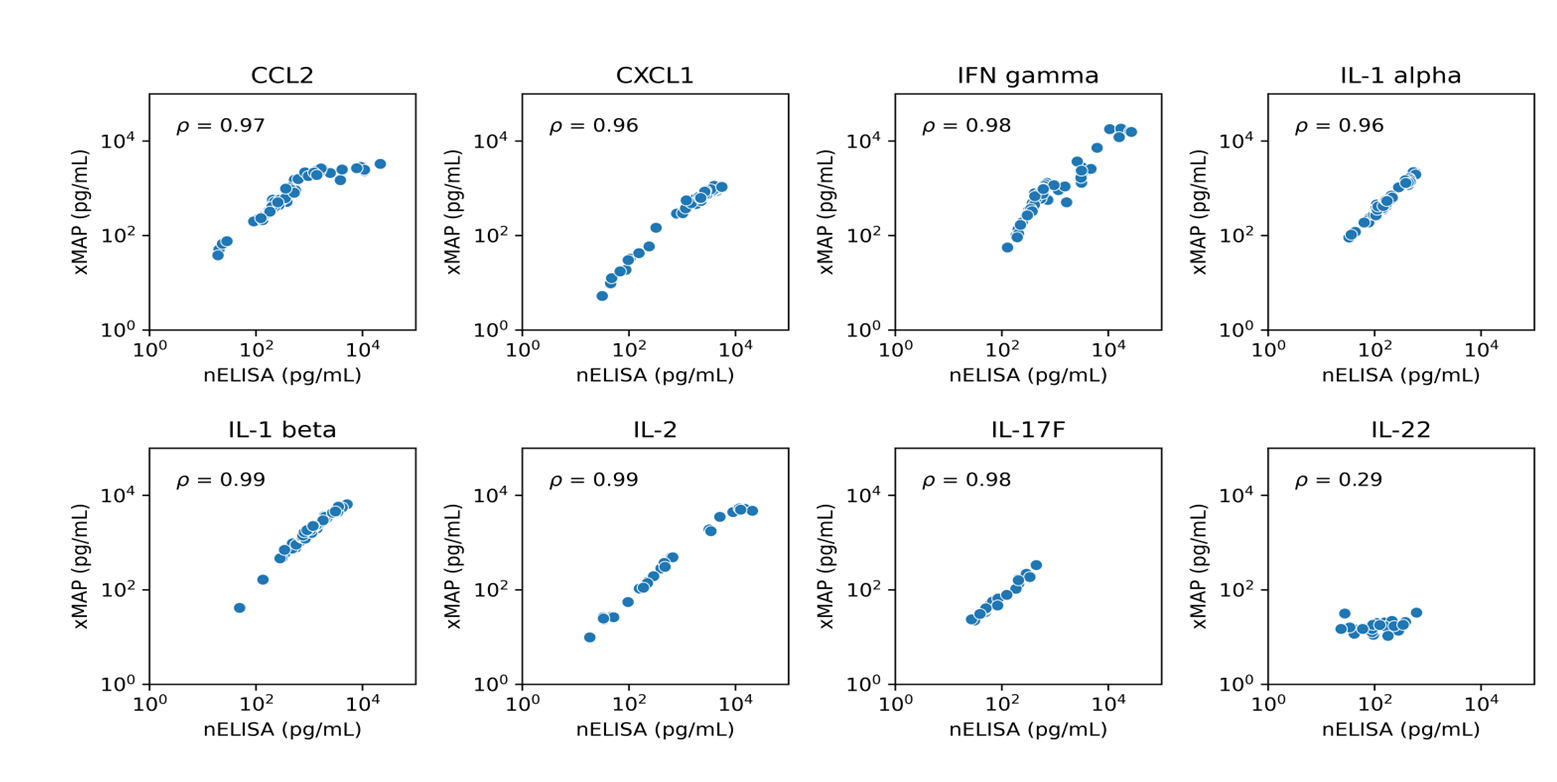

**Suppl. Fig. 8:** Correlation between nELISA and xMAP. Cytokine levels in cell culture supernatants from stimulated PBMCs profiled with the nELISA 191-plex and a 48-plex panel based on the xMAP platform. Shown are the detected protein levels for a subset of sensors shared by both platforms and yielding detectable protein concentrations, with associated Spearman correlations. Some of the lower Spearman correlations are explainable. Notably, IL-22 was observed to saturate in xMAP, whereas it was in the quantifiable range with nELISA.

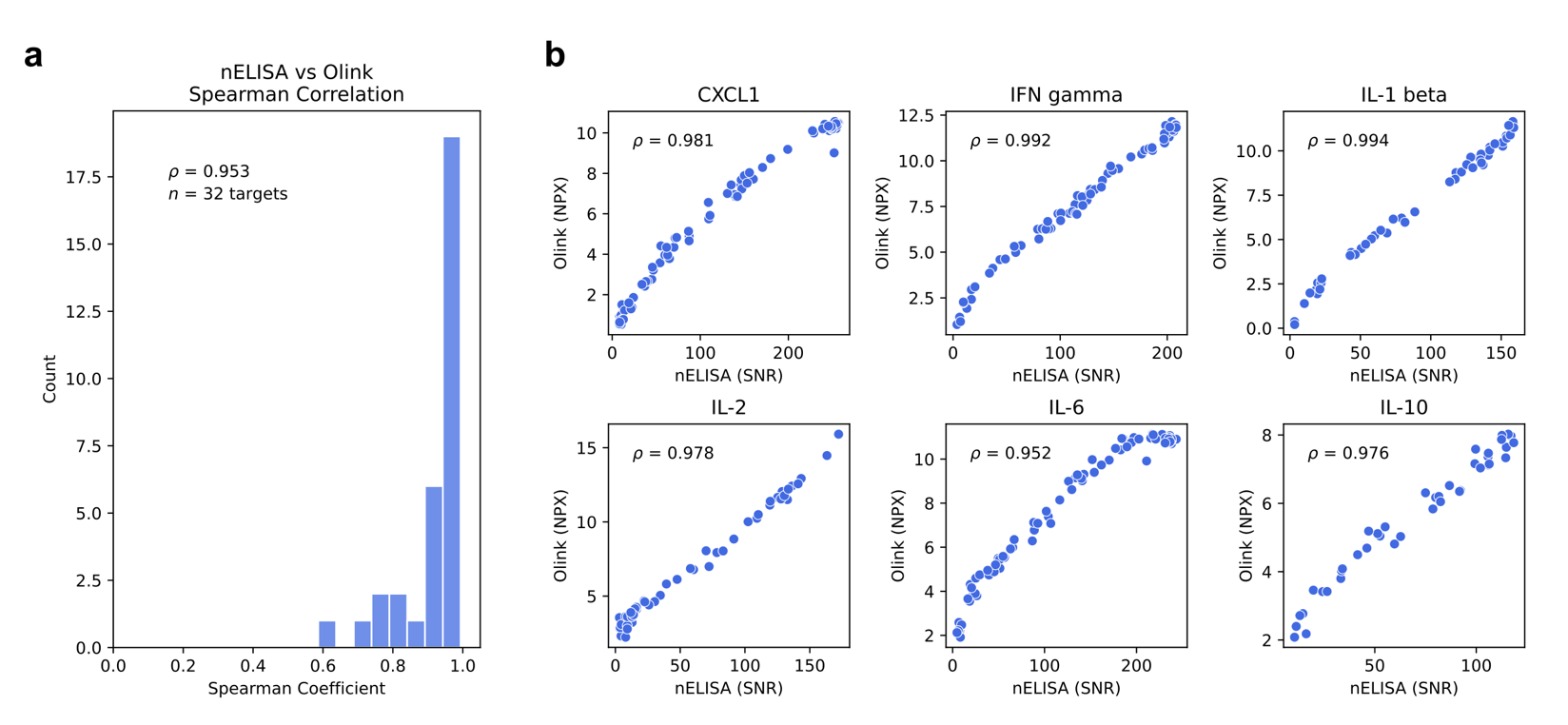

**Suppl. Fig. 9: Correlation Between nELISA and Olink in Cell Supernatant Samples.** PBMCs from four donors were treated with four concentrations of each stimulus (LPS, ConA, PolyIC, PMA/I) or left unstimulated for 24 hours. Cytokine levels were quantified using the nELISA 191-plex and the Olink Explore 384 Inflammation panel. Of the 85 shared targets, 36 had detectable expression above the limit of detection on both platforms, and 31 had no expression on either platform (in >95% of data points) and 18 had expression on one platform only. (a) Histogram of Spearman correlation coefficients for all detected targets, limited to high-confidence correlations (p-value > 0.05). The median correlation exceeded 95%, demonstrating strong concordance between platforms. (b) Cross-correlation plots for selected targets, comparing nELISA (SNR) and Olink (NPX) values for data above the detection limit on each platform. Notably, IL-6 exhibited saturation in Olink, whereas nELISA measurements remained within the quantifiable range.

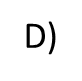

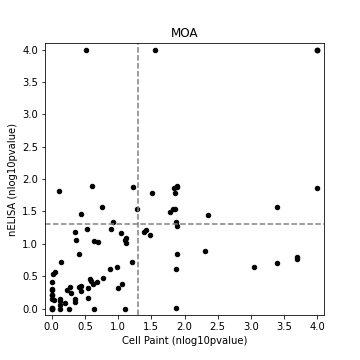

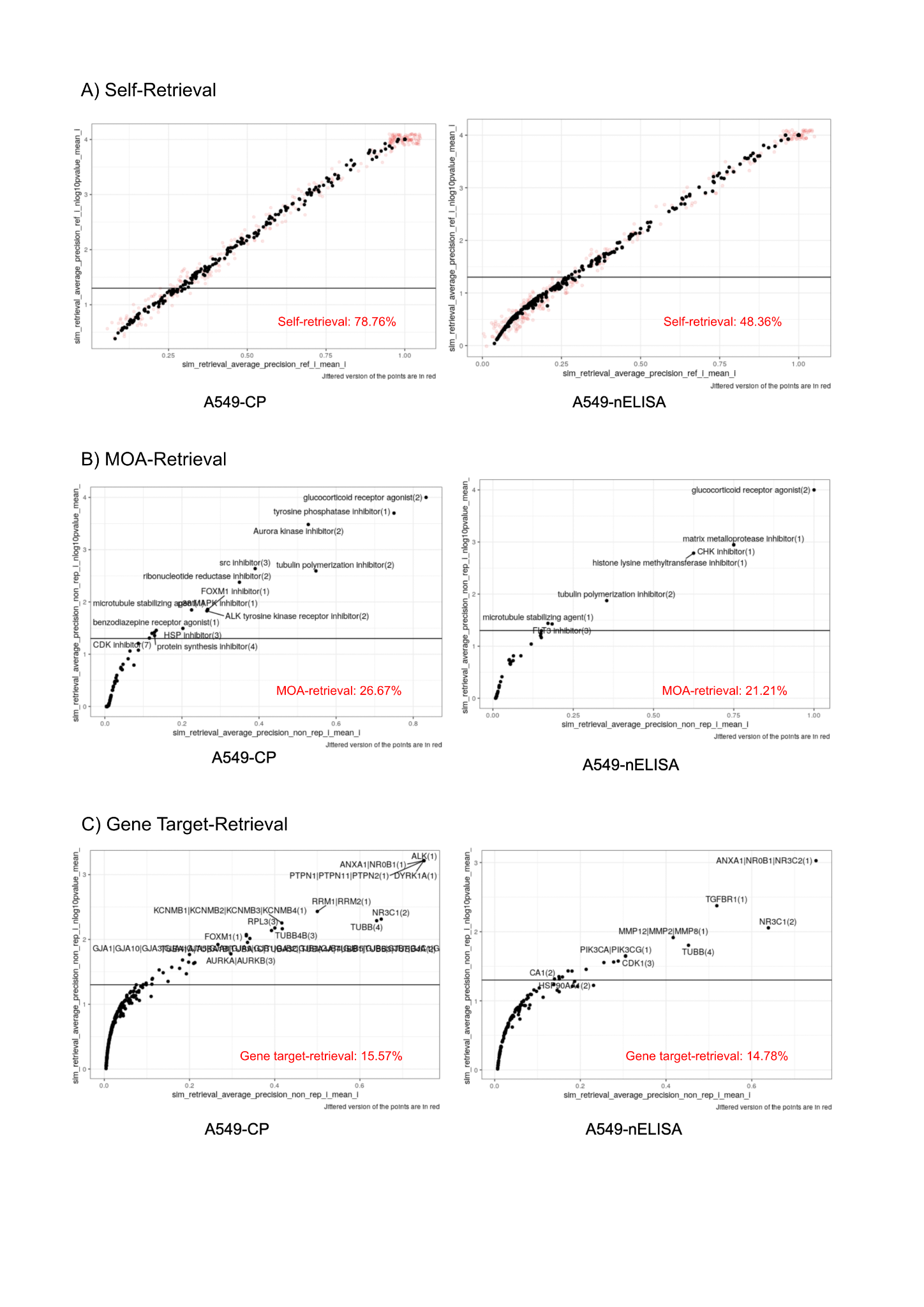

#### Suppl. Fig. 10: nELISA and Cell Painting reveal complementary biology. Distribution of compound phenotypic activity, as measured by mean average precision (mAP) scores for replicate wells of A549 cells treated with a small molecule library; the horizontal line indicates the significance threshold (p=0.05) for compounds that could reliably retrieve their replicate samples (self-retrieval) against DMSO control-treated A549 cells, using Cell Painting (left) and nELISA (right) readouts. A jittered version of the data is shown in red to show the density of overlapping data points. (B) Distribution of compound mAP scores for compounds with shared MOAs; the horizontal line indicates the significance threshold (p=0.05) for compounds that could reliably retrieve compounds annotated with the same MOA (MOA retrieval, a measure of phenotypic consistency). (C) Distribution of compound mAP scores for compounds with shared gene targets; the horizontal line indicates significance threshold (p=0.05) for compounds that could reliably retrieve compounds annotated with the same gene target (gene target retrieval, a measure of phenotypic consistency). D) Scatterplot of MOA-retrieval on nELISA and Cell Painting; vertical and horizontal lines indicate significance threshold. E) Scatterplot of gene target-retrieval on nELISA and Cell Painting; vertical and horizontal lines indicate significance threshold. Note: These convenience samples were suboptimal for secretome profiling in three ways: first, the experiment used A549 cells, which are relatively non-secretory, second, the cells were not subjected to any stimulatory conditions, and third, the goal of the experiment was to identify responses to a wide variety of drugs, impacting various diseases and pathways, with the vast majority unrelated to the processes targeted by the 191-plex protein panel.

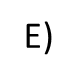

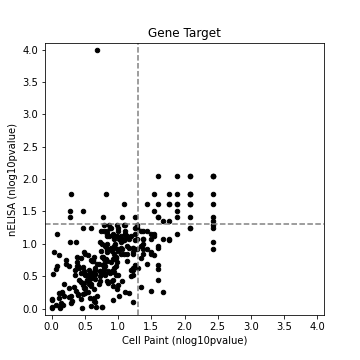

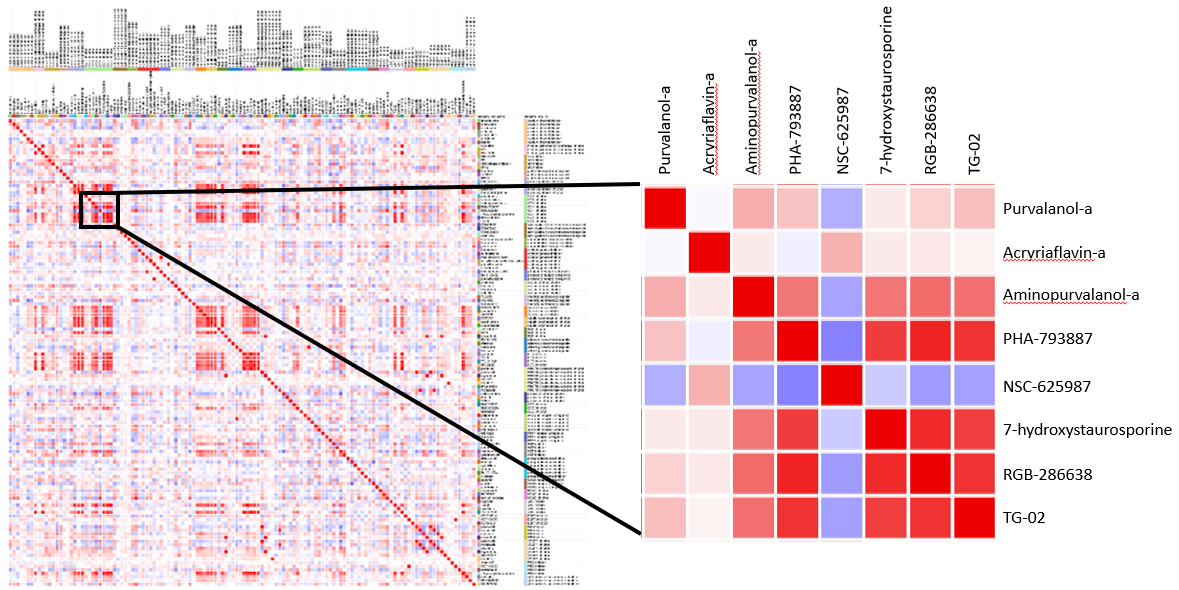

#### Suppl. Fig. 11: nELISA reveals phenotypic differences within mechanisms of action categories.

(left) Correlation of cosine similarity scores across compounds. Compounds are grouped by MOA, resulting in red squares around the diagonal where compounds within an MOA have high cosine similarity scores. Only MOAs with 3 or more compounds are included. Blue indicates negative similarity scores, red indicates positive similarity scores. (right) Magnification of the MOA category “CDK inhibitors”, which includes 8 compounds. MOA retrieval of nELISA data only led to the prediction of 1 additional compound when compared with Cell Paint (Suppl. Fig 5b), however, pairwise comparisons uncovered that at least 5 of these compounds had high cosine similarity scores, with a single anti-correlated compound (NSC-625987). All the CDK inhibitors other than NSC-625987 target multiple CDKs and other kinases, whereas NSC-625987 is specific to CDK4, indicating that nELISA results could distinguish compounds with similar mechanisms of action.

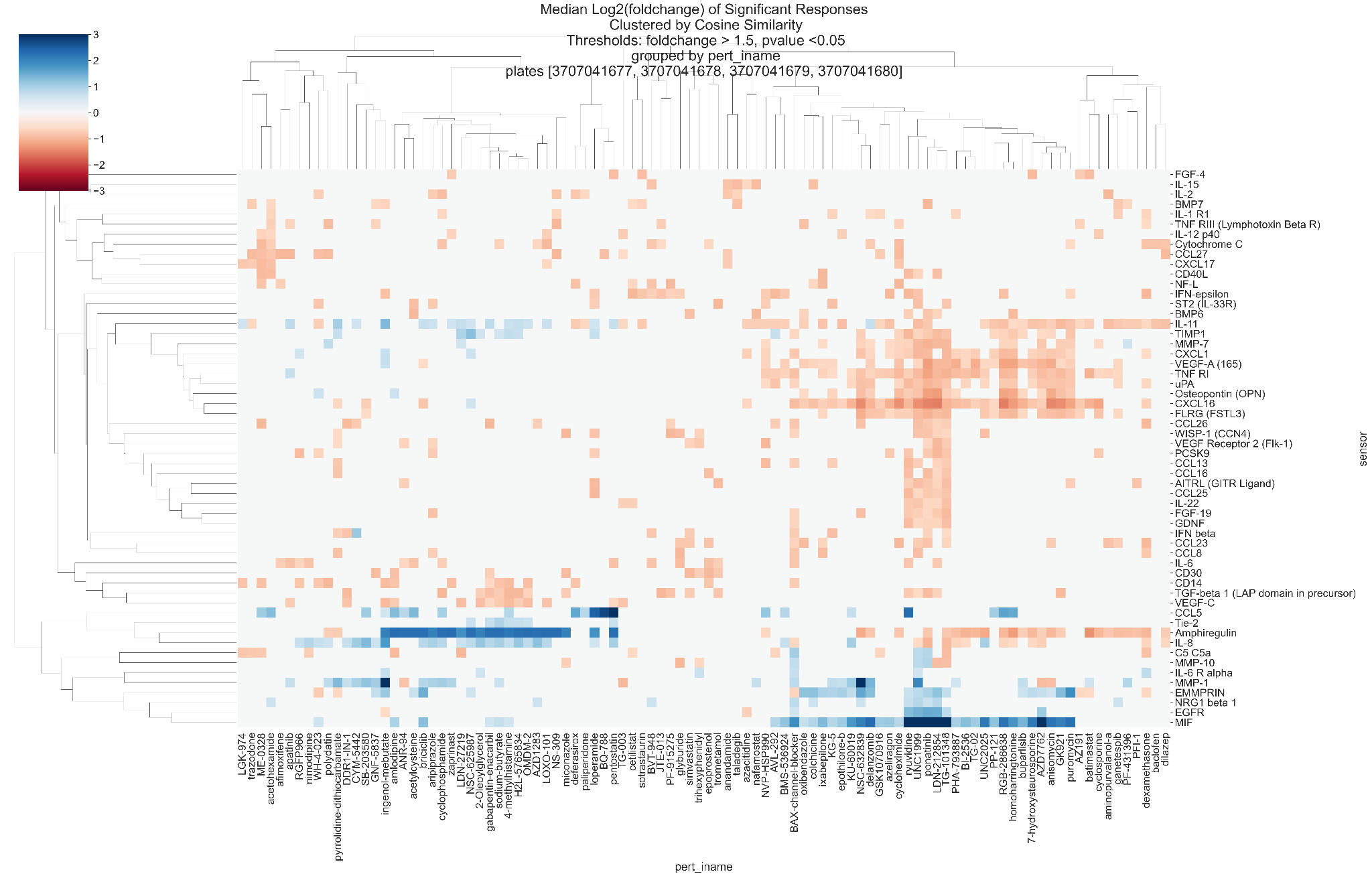

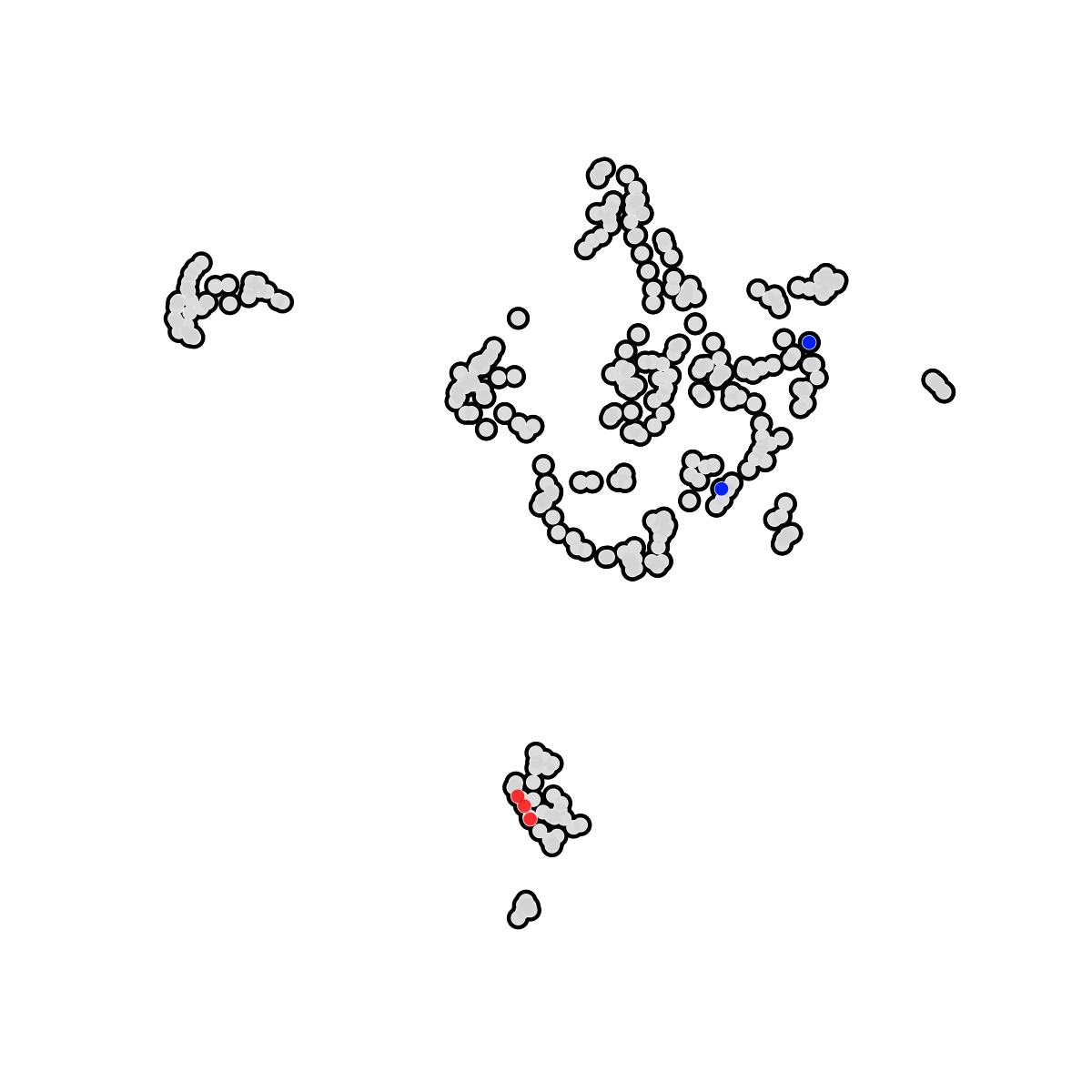

#### Suppl. Fig. 12: A549 phenotypic screening. (top) Heatmap dendrogram of significant changes in cytokine expression levels in response to treatment with indicated compounds. (bottom) UMAP clustering of compounds with shared effects on A549 secretomes; CHK inhibitors and the downstream Aurora B/C kinase inhibitor GSK1070916 are identified in red; pan-Aurora kinase inhibitors danusertib and AMG900 are colored blue.

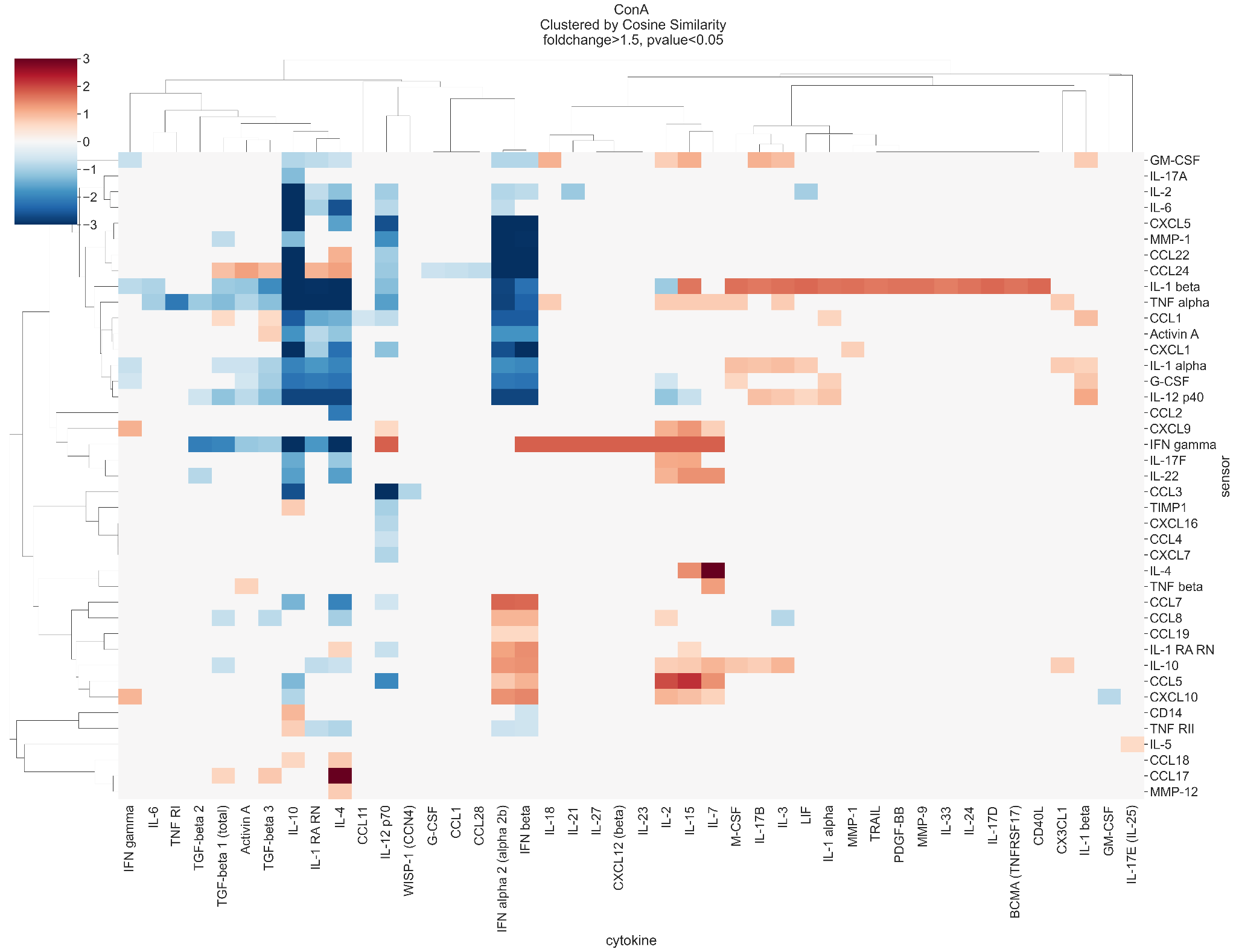

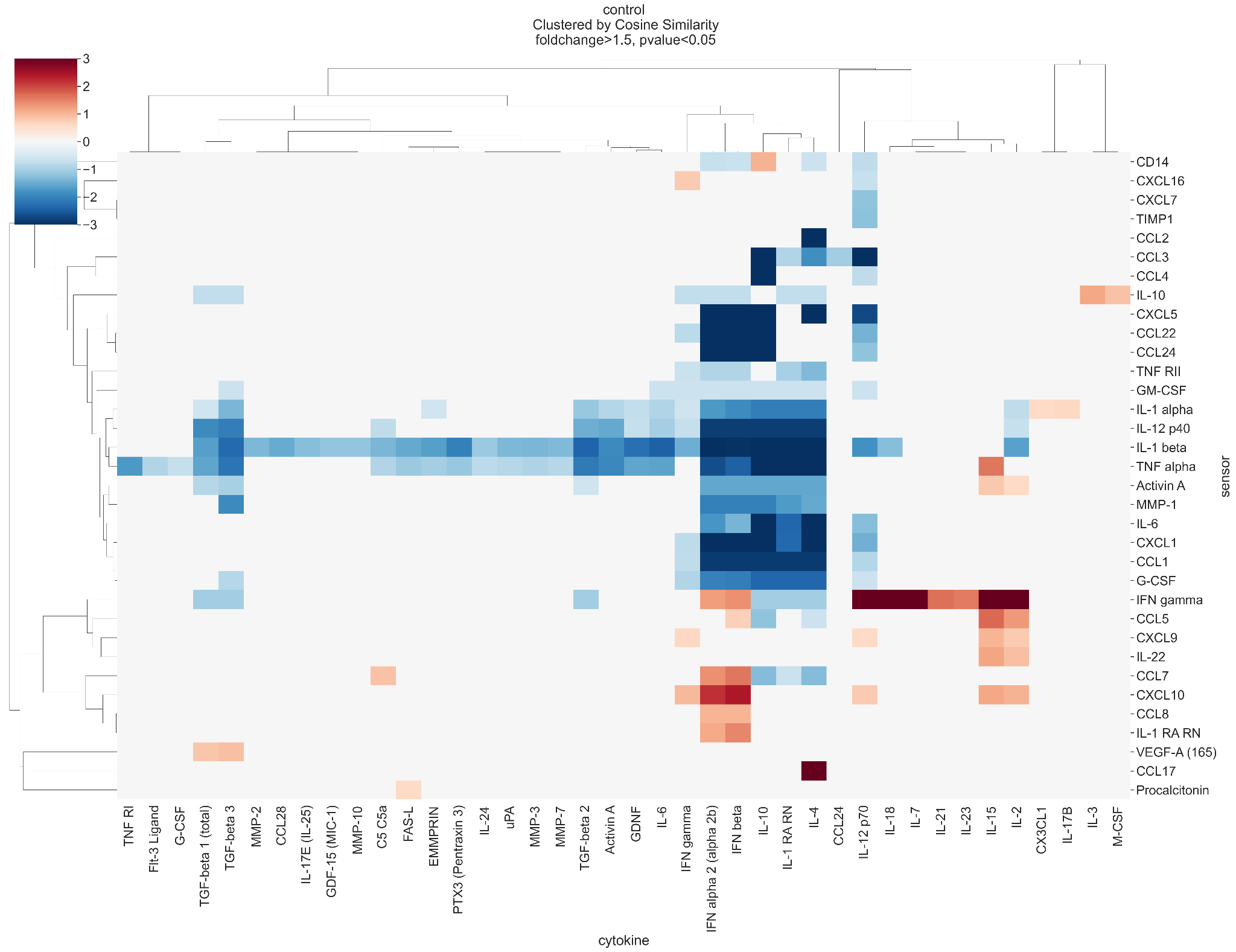

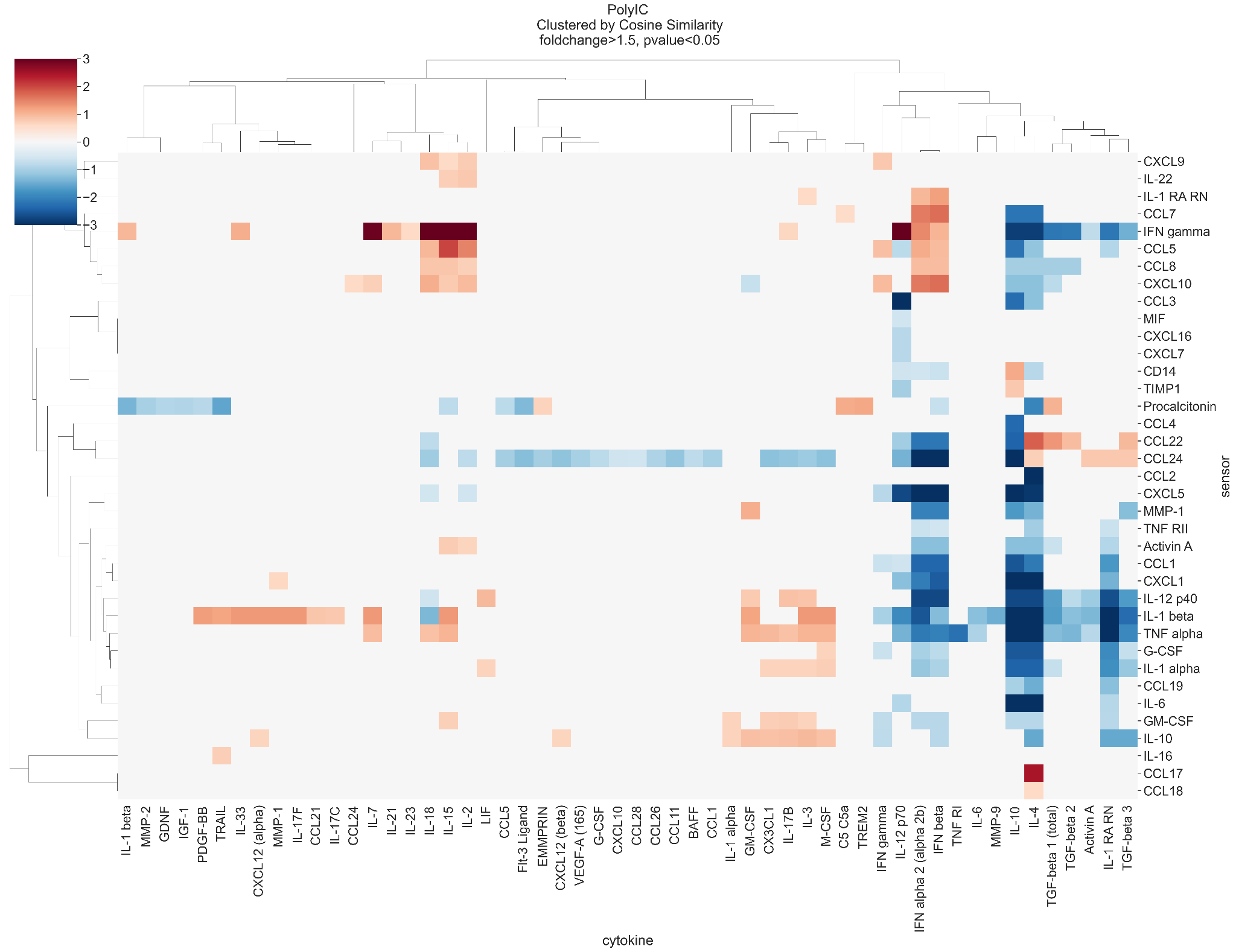

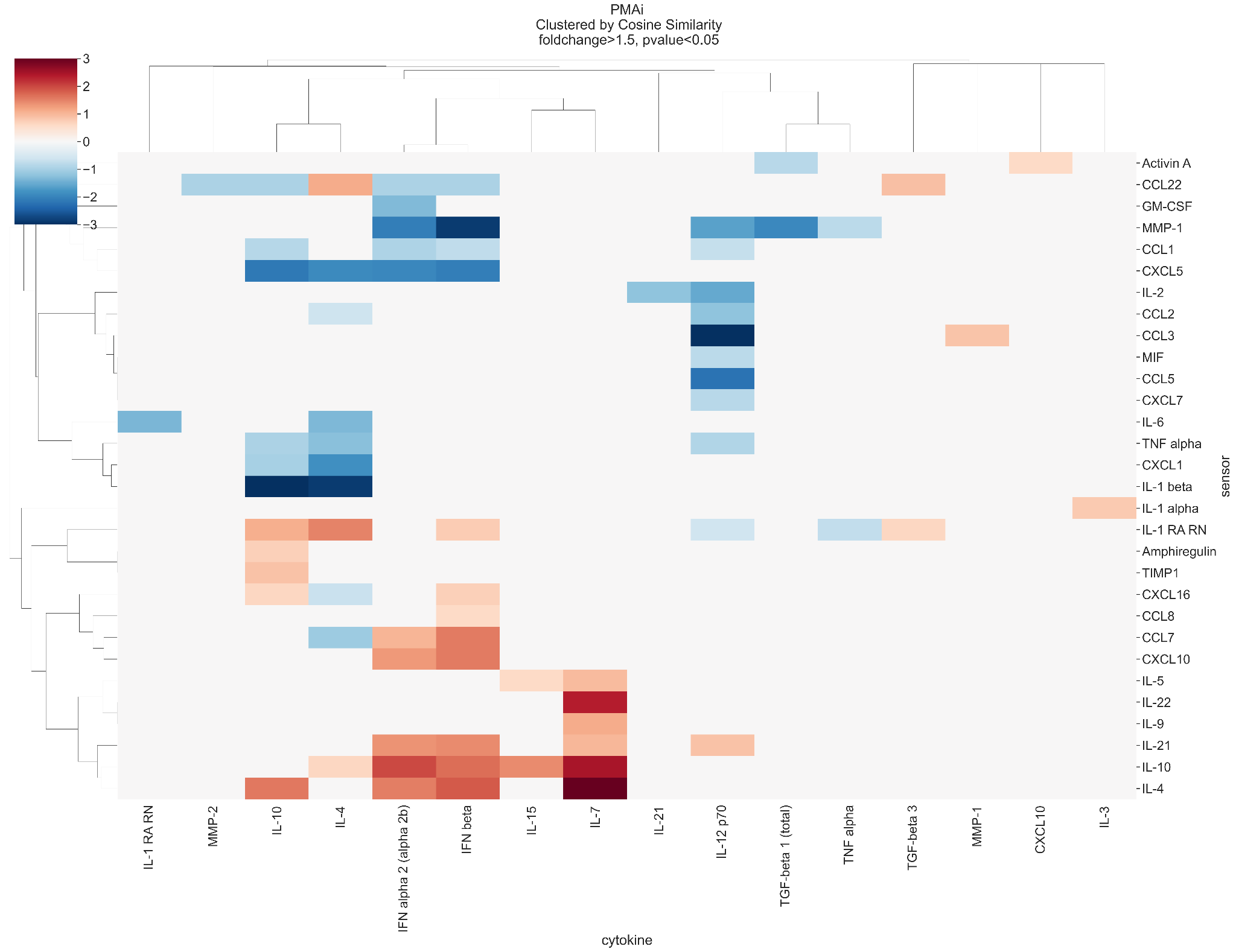

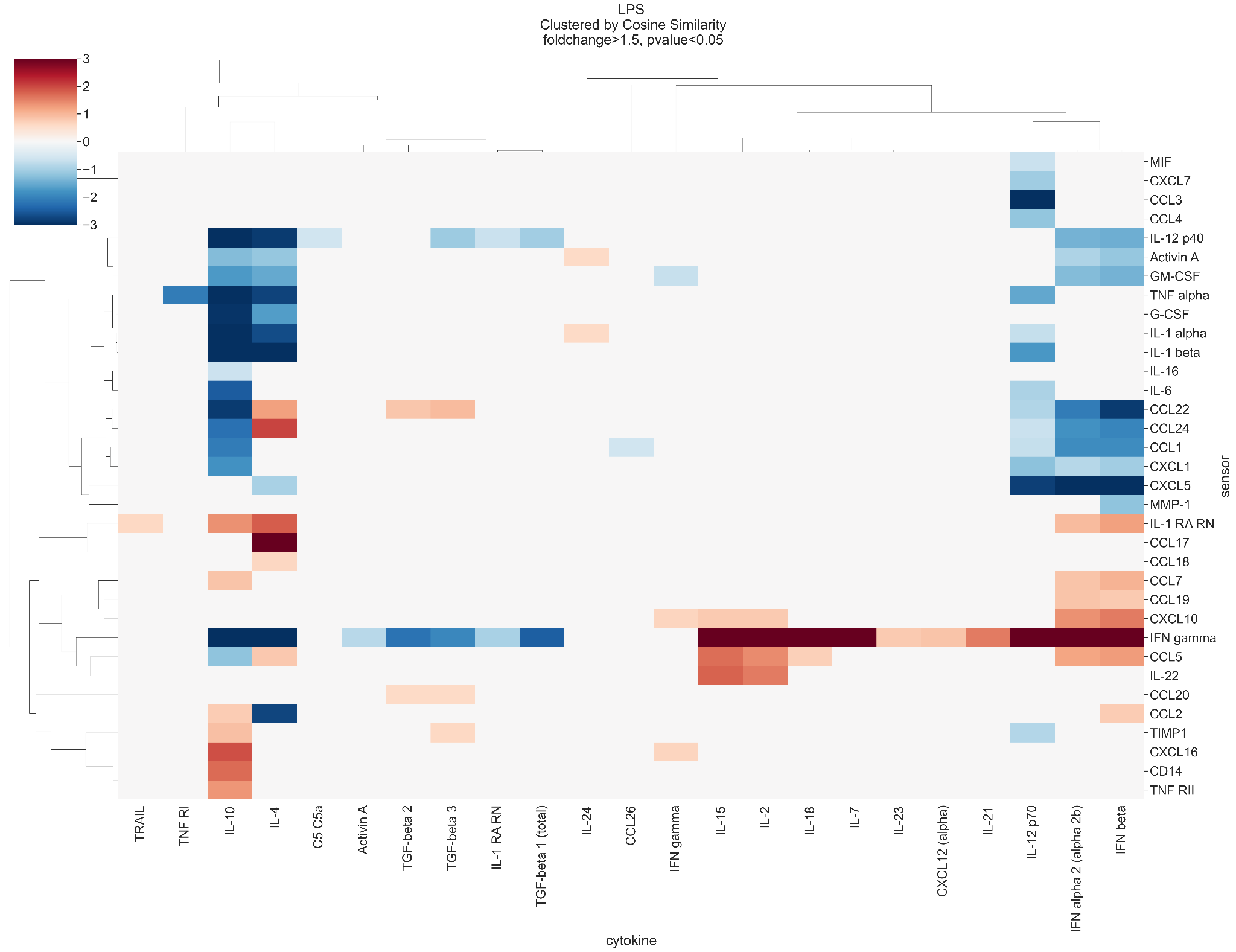

#### Suppl. Fig. 13: Heatmap dendrograms of cytokine interactions in each stimulus condition. PBMCs were treated with indicated stimuli (ConA, Control, LPS, PMA/i or PolyIC; a separate heatmap is shown for each), and a library of 80 recombinant cytokines; shown are the significant fold changes in expression of each sensor in response to each recombinant cytokine.

### **Supplemental Calculation 1.** Antibody concentration at the bead surface and in solution after oligo displacement and release.

#### 1. Surface Area of a Single Bead

Assume each bead is a sphere with a diameter of 3 µm.
The surface area (A) of a sphere is given by: A = 4πr²
A_bead_ = 4π(1.5 µm)² ≈ 28.3 µm² per bead

#### 2. Approximate Number of Antibodies on a Single Bead

Assume an IgG has dimensions of approximately 14.5 nm × 8.5 nm x 4 nm. [1]
Assume 20% coverage of the surface with detection antibody (capture oligo represents 20% of total DNA on bead).

The area occupied by one antibody is estimated by the average of each face:

A_IgG_ = (A_face1_ + A_face2_ + A_face3_) / 3

A_IgG_ = ((14.5 nm × 8.5 nm) + (14.5 nm × 4 nm) + (8.5 nm × 4 nm)) / 3
A_IgG_ = 71.7 nm^2^ = 7.17 × 10^-5^ µm^2^

The number of detection antibodies on the bead surface at 20% coverage is:
N_IgG_ = 0.2 × (A_bead_ / A_IgG_)
N_IgG_ = 0.2 × (28.3 µm²) / (7.17 × 10^-5^ µm^2^)
N_IgG_ = 78,940

#### 3. Approximate Local Antibody Concentration at the Bead Surface

Approximate the local concentration by assuming the antibodies are confined within a thin shell around the bead, roughly 10 nm (0.01 µm) thick.

The volume of this shell around a single bead is approximated as:
V_shell_ ≈ 4/3π(r_total_^3^ - r_bead_^3^)
V_shell_ ≈ 4/3π((3.01 µm)^3^ – (3.00 µm)^3^)
V_shell_ ≈ 1.1 µm^3^ ≈ 1.1 × 10^-15^ L

The molar concentration of detection antibodies in this shell is given by:
C_surface_ = N_IgG_ / (N_A_ × V_shell_) where N_A_ is Avogadro’s number.
C_surface_ = (78,940)/ ((6.022 x 10^23^) × (1.1 x 10^-15^ L))
C_surface_ = 1.19 × 10^-4^ M = 119 µM

#### 4. Approximate Antibody Concentration in Solution After Release

Assume 50 nELISA beads per target in each sample well.
Assume detection antibody release efficiency is 100%.
Assume antibodies are released into 100 µL.

Total number of antibodies (N_total_) released:
N_total_ = 50 beads × N_IgG_
N_total_ = 50 × (78,940) = 3.9 × 10^6^

The total solution volume (V_total_) is 100 µL = 100 × 10^-6^ L.
The final concentration in solution (C_solution_) is given by:
C_solution_ = N_total_ / (N_A_ × V_total_)
C_solution_ = (3.9 × 10^6^) / ((6.022 x 10^23^) × (100 × 10^-6^ L))
C_solution_ = 6.55 × 10^-14^ M = 65 fM

**Supplemental Table 1:** Comparison of cost and throughput for protein-profiling technologies.

| **Specification** | | **nELISA** | | **xMAP** | | **Olink** | |
| --- | --- | --- | --- | --- | --- | --- | --- |
|  | | 30-plex | 191-plex | 34-plex ^[1]^ | 80-plex ^[2]^ | Target 48^[3]^ | Target 96^[4]^ |
| Throughput | Number of samples/day | 1,440 ^[4]^ | 1,440 ^[4]^ | 80 | 76 | 120 | 264 |
|  | Number of samples/week | 10,080 | 10,080 | 560 | 532 | 840 | 1,848 |
|  | Plate format | 96 or 384 | 96 or 384 | 96 | 96 | 96 | 96 |
| Cost | Per sample pricing | <$20 USD | <$40.00 USD | $151.25 USD | $293.00 USD | $122.50 USD | $72.72 USD |
|  | Cost per datapoint | <$0.67 USD | <$0.21 USD | $4.44 USD | $3.66 USD | $2.72 USD | $0.83 USD |
|  | Proprietary Equipment | No | No | Yes | Yes | Yes | Yes |
| Assays | Flexibility | All targets compatible in custom panels. | | Pre-set panels with limited compatibility combining panels. | | Pre-set panels with limited compatibility combining panels. | |
|  | Quantitative | Yes | | Yes | | Yes | No |

[1] [ProcartaPlex™ Human Immune Response Panel, 80plex.](https://www.thermofisher.com/order/catalog/product/EPX800-10080-901?SID=srch-srp-EPX800-10080-901)

[2] [ProcartaPlex™ Human Cytokine & Chemokine Panel 1A, 34plex](https://www.thermofisher.com/order/catalog/product/EPXR340-12167-901)
[3] Olink Target-48: 40 samples per 96 wellplate with 45 assays (quantitative).

[4] Olink Target-96: 88 samples per 96 wellplate with 92 assays (non-quantitative).

[5] nELISA profiling 1440 samples per day per unit cell (see supplementary table 2 for scalability analysis).
