## Supplemental Protocol for "nELISA: A high-throughput, high-plex platform enables quantitative profiling of the inflammatory secretome"

**nELISA Protein Profiling Protocol**

| **Buffer** | **Composition** |
| --- | --- |
| Culture Media | RPMI, 10% HI-FBS |
| Wash Buffer | PBST 0.1%, 4 mM EDTA |
| Displacement Buffer | PBS (1x), 1% BSA, 4 mM EDTA, 900 mM NaCl, 0.02 µM displacement oligo |
| Lysate Buffer | M-PER (Thermo Fisher Scientific), 1x protease/phosphatase inhibitor (Thermo Fisher Scientific)  20mM EDTA, 20mM EGTA, 100 ug/mL salmon sperm DNA |

| **Stage** | **Steps** | **Approx. Time** |
| --- | --- | --- |
| Sample Preparation | Samples are thawed for 1 hour on ice while covered. Once thawed, sample plates are shaken at 1800 rpm for 2 minutes (Bioshake XP shaker; Bulldog Bio) and centrifuged at 500 × g for 15 seconds (Sorvall Legend XFR Refrigerated Centrifuge; Thermo Scientific).  Samples are diluted into 384 wellplates of culture media at 2-fold and 50-fold using a liquid handler (Viaflo; Integra), followed by mixing at 1800 rpm for 2 minutes and centrifugation at 3800 × g for 5 minutes.  For cell lysates, sample preparation is performed as above using lysate buffer in place of culture media. | 1 hour + 30 min |
| nELISA Assay | Wash buffer and displacement buffer are prepared fresh on the day of the assay.  Samples are transferred to assay plates containing nELISA beads via liquid handler (Viaflo; Integra). Assay incubation is carried out at room temperature for 3 hours at 2400 rpm Bioshake XP shaker; Bulldog Bio).  After incubation, target-bound nELISA beads undergo four washes. For each wash cycle, nELISA beads are pelleted by centrifugation (500 × g, 30 seconds, 4°C), the wash buffer is exchanged using a plate washer (MultifloFX; Biotek), plates are washed at 2200 rpm for 30 seconds.  With the final wash, 50 µL of wash buffer is left in each well. An equal volume (50 µL) of displacement buffer is added and plates are incubated at room temperature for 30 minutes at 2200 rpm.  Following the displacement reaction, nELISA beads undergo an additional four wash cycles (as described above). With the final wash, 18 µL of wash buffer remains in each well.  nELISA beads are resuspended by shaking at 2400 rpm and measured by high-throughput flow cytometry (ZE5 cell analyzer; Bio-Rad). Acquisition is performed on high-throughput mode following calibration with rainbow beads (Sphero Rainbow Calibration Particles; BD Biosciences). A cleaning cycle (bleach, cleaner, water) is performed between each plate readout. | 3 hours (sample incubation)  + 3 hours (washes, DO, and readout) |
